# Enhanced metabolic detoxification is associated with fluroxypyr resistance in *Bassia scoparia*

**DOI:** 10.1101/2023.08.29.554743

**Authors:** Olivia E. Todd, Eric L. Patterson, Eric P. Westra, Scott J. Nissen, André Lucas Simões Araujo, William B. Kramer, Franck E. Dayan, Todd A. Gaines

**Affiliations:** United States Department of Agriculture – Agriculture Research Service, Fort Collins, CO 80525, USA; Department of Agricultural Biology, Colorado State University, Fort Collins, CO 80523, USA; Department of Plant, Soil, and Microbial Sciences, Michigan State University, East Lansing, MI 48824, USA; Department of Plants, Soils & Climate, Utah State University, Logan, UT 84322, USA

**Keywords:** Herbicide Resistance, NTSR, Synthetic Auxin

## Abstract

Auxin-mimic herbicides chemically mimic the phytohormone indole-3-acetic-acid (IAA). Within the auxin-mimic herbicide class, the herbicide fluroxypyr has been extensively used to control an agronomically problematic Great Plains tumbleweed, kochia (*Bassia scoparia*). A 2014 field survey for herbicide resistance in kochia populations across Colorado identified a putative fluroxypyr resistant population that was assessed for response to five different herbicides representing four different herbicide modes of action. These included fluroxypyr and dicamba (auxin-mimics), atrazine (photosystem II inhibitor), glyphosate (EPSPS inhibitor), and chlorsulfuron (acetolactate synthase inhibitor). The greenhouse screen identified that this kochia population was resistant to fluroxypyr and chlorsulfuron, but sensitive to glyphosate, atrazine, and dicamba. This population was designated Flur-R. Subsequent dose response studies determined that 75% of the Flur-R population survived 628 g ae ha^-1^ of fluroxypyr (4X the label application rate in wheat fallow, which is 157 g ae ha^-1^ at 1X). Flur-R was 40 times more resistant to fluroxypyr than a susceptible population (J01-S) collected from the same field survey (LD_50_ 720 and 20 g ae ha^-1^, respectively). Auxin-responsive gene expression increased following fluroxypyr treatment in Flur-R, J01-S, and in a dicamba-resistant, fluroxypyr-susceptible line 9425 in an RNA-sequencing experiment. In Flur-R, several transcripts with molecular functions for conjugation and transport were constitutively higher expressed, such as glutathione S-transferases (GSTs), UDP-glucosyl transferase (GT), and ATP binding cassette transporters (ABC transporters). After analyzing metabolic profiles over time, both Flur-R and J01-S rapidly converted [^14^C]-fluroxypyr ester, the herbicide formulation applied to plants, to [^14^C]-fluroxypyr acid, the biologically active form of the herbicide, and three unknown metabolites. Formation and flux of these metabolites was faster in Flur-R than J01-S, reducing the concentration of phytotoxic fluroxypyr acid. One unique metabolite was present in Flur-R that was not present in the J01-S metabolic profile. Gene sequence variant analysis specifically for auxin receptor and signaling proteins revealed the absence of non-synonymous mutations affecting auxin signaling and binding in candidate auxin target site genes, further supporting our hypothesis that non-target site metabolic degradation is contributing to fluroxypyr resistance in Flur-R.

**Significance Statement:** Herbicide resistance is an ever-present issue in weeds of cropping and rangeland systems. By understanding genetic mechanisms of resistance in individual cases of herbicide resistance, we can extrapolate important information such as how quickly resistance to a specific herbicide can spread. Every characterized herbicide resistance mechanism contributes to a working database used to address herbicide resistance in an agricultural or open-space setting. Knowing the exact mechanism of resistance helps researchers and industry members understand why herbicide applications are failing, and if resistant plants can still be controlled with other herbicide modes of action. In kochia line Flur-R, there is strong evidence to support a non-target site resistance mechanism, specifically herbicide degradation via increased enzymatic activity. Increased fluroxypyr degradation represents a novel resistance mechanism to fluroxypyr in the weed *Bassia scoparia*.

## 1. Introduction

Kochia (*Bassia scoparia*) is an invasive, annual, broadleaf tumbleweed that is problematic in agronomic settings, open spaces, and rangeland in the U.S., specifically across the Great Plains. Herbicides, chemicals used to control unwanted plants, are the most prescribed method for kochia control in the U.S. Herbicide resistance in kochia is widespread, with resistance reported to multiple modes of action including ALS inhibitors (Group 2), glyphosate (Group 9), auxin-mimics (Group 4), and atrazine (Group 5), as well as cross-resistance to more than one Group 4 auxin-mimic herbicide (Geddes et al. 2021a; Kumar et al. 2019a; Kumar et al. 2019b). Furthermore, multiple resistance is increasingly common, where one population that is simultaneously resistant to more than one mode of action (Varanasi et al. 2015). The prevalence of glyphosate, atrazine, and ALS inhibitor resistant kochia has resulted in increased use of auxin-mimic herbicides for kochia management, mainly dicamba and fluroxypyr, in no-till fallow, wheat, and corn in the Great Plains region (Kumar et al. 2019b). While auxin-mimics have been used for more than 70 years, the evolution of resistance to this mode of action has lagged behind other herbicide modes of action (Busi et al. 2018). Despite this lag, recent evidence suggests that auxin-mimic resistance and multiple resistance in kochia is increasing (Geddes et al. 2022). Nine reports of auxin-mimic resistance across six U.S. states and two Canadian provinces have described resistance in kochia populations as either resistant to dicamba alone, or resistant to both dicamba and fluroxypyr (Geddes et al. 2021b; Heap 2021; Jha et al. 2015; Kumar et al. 2019a). These herbicides mimic the phytohormone indole-3-acetic-acid (IAA) because they are chemically similar and induce auxin response gene transcription following application in weeds (Grossmann 2010; McCauley et al. 2020; Pettinga et al. 2018; Xu et al. 2022) (Figure 1).

**Figure 1:**
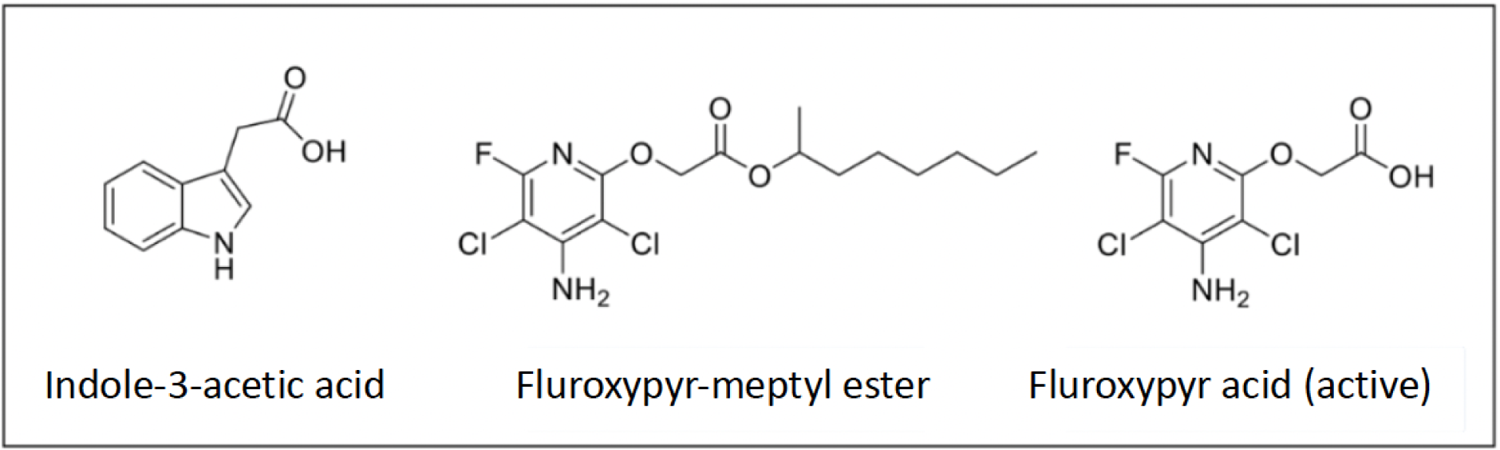
Chemical structure of indole-3-acetic acid (IAA), fluroxypyr-meptyl ester (fluroxypyr-ester) included in the formulated commercial products, and fluroxypyr-acid, the biologically active form of the herbicide. Deesterification of fluroxypyr-meptyl ester frees the carboxyl group shown in fluroxypyr-acid, which plays a key role in plant perception related to the auxin signaling pathway in relation to the ring structure found throughout auxin-mimic herbicide chemistry.

IAA is an auxin plant growth hormone that is responsible for gravitropism and response to light stimuli; however, it impacts several other growth phenotypes as well (Zhao 2010). While auxins are involved in many cellular processes and signaling with other phytohormones, their function can be understood at the cellular level to primarily coordinate cell elongation (Perrot-Rechenmann 2010). In plants, auxin homeostasis is tightly regulated through a suite of biosynthesis pathways, cellular transport, feedback inhibition, oxidation and conjugation (Rosquete et al. 2012). When IAA reaches high levels in the plant, polar auxin carriers such as Pin-formed (PIN) efflux transporters, ATP binding cassettes (ABC class B) and Auxin resistant-1/like AUX1s influx carriers (AUX/LAX) help maintain IAA homeostasis and gradients (Cho and Cho 2013). Because the auxin-mimic herbicide fluroxypyr is chemically similar to and behaves like IAA, it is hypothesized that PIN, ABCs and AUX/LAX are able to direct the flow of fluroxypyr throughout the plant. In addition, fluroxypyr is a weak acid that can translocate in the plant based on its pKa and log Kow. Fluroxypyr also binds to Auxin signaling F-box 5, a member of the Transport inhibitor response1/Auxin signaling F-box (TIR1/AFB) receptor family (Lee et al. 2014). When applied, fluroxypyr and IAA act to stabilize the complex formed between AFB5 and the auxin dependent transcriptional regulator Indole-3-acetic acid inducible (Aux/IAA) proteins. Upon creation of this coreceptor-ligand complex, Aux/IAA proteins are ubiquitinated, degraded, and no longer negatively regulate Auxin Response Factors (ARF). These ARF proteins are seated on the Auxin Response Element (AuxRE) in auxin mediated gene promotors (Teale et al. 2006).

In Arabidopsis, plants treated with IAA or auxin-mimic herbicide (2,4-D) showed expression of early response genes such as Aux/IAAs and small auxin-up RNAs (SAURs). Other auxin induced genes included 1-aminocyclopropane-1 synthase (ACS), the first committed step in ethylene production and GH3, an auxin homeostasis gene. These genes were transcribed within minutes of high auxin perception (Guilfoyle 1999; Paponov et al. 2008; Raghavan et al. 2005). Many other phytohormone responses are also regulated by auxin perception, such as Cytokinin oxidase (CXK6), brassinosteroid biosynthesis gene BAS1, and several gibberellin related genes. Regulation of multiple hormone related genes suggests that the relationship between phytohormones and auxin response is complex (Paponov et al. 2008). When the auxin-mimic herbicide fluroxypyr is applied to a plant, the resulting phenotypic response is stem-twisting, thickening, and lack of new growth at the meristem.

Herbicide resistance is categorized as either target site or non-target site (Gaines et al. 2020). Target site resistance is defined as a change (either in conformation or in expression) in a herbicide target protein. These changes often decrease herbicide binding affinity or in the case of overexpression, make it so the entire target protein pool is unable to be entirely inhibited (Murphy and Tranel 2019). LeClere et al. (2018) reported resistance to the auxin-mimic herbicide dicamba in kochia was due to the target-site mutation Gly127Asn in *IAA16*, which affects the formation of the coreceptor-ligand complex. More recently, Figueiredo et al. (2022b) characterized a 27-nucleotide deletion in the degron tail region of the gene encoding Aux/IAA2 that confers 2,4-D and dicamba resistance in Indian hedge mustard (*Sisymbrium orientale*). This deletion also affects formation and stability of the co-receptor-ligand complex. Non-target site resistance is broadly recognized as all other methods unrelated to target site resistance and is often exemplified by metabolic detoxification of an herbicide, herbicide sequestration, or a variant in a metabolism catalyzing enzyme which may have a downstream effect by reducing the efficacy of the herbicide (Delye 2013). Reduced translocation of 2,4-D via auxin transport proteins was reported by Goggin et al. (2019) in wild radish (*Raphanus raphanistrum*). A 2,4-D resistant population of waterhemp (*Amaranthus tuberculatus*) rapidly metabolized 2,4-D via CYP450 5-OH hydroxylation and subsequent amino acid and sugar conjugation reactions, which produced less phytotoxic metabolites that lost auxin signaling activity (Figueiredo et al. 2022a; Figueiredo et al. 2018). With both target site and non-target site resistance mechanisms described for auxin-mimic resistance (Todd et al. 2020), both possibilities are investigated in this study.

Our research objectives were to (1) **distinguish the application rate at which the fluroxypyr resistant kochia line (Flur-R) is resistant using a herbicide dose response,** (2) **determine whether fluroxypyr resistant kochia has any differences in absorption, translocation, or metabolism of fluroxypyr**, and (3) **identify potential candidate genes that may contribute to fluroxypyr resistance in Flur-R using RNA sequencing and alignment to the kochia genome assembly** (Hall et al. 2023).

In our work, a kochia population that was resistant to fluroxypyr converted fluroxypyr-ester into fluroxypyr-acid and subsequent metabolites at a faster rate than a susceptible line. Furthermore, the resistant line produced a metabolite that was not detected in the susceptible line. The results from an RNA-seq fluroxypyr-induced differential expression analysis show increased transcript expression of cellular transporters, cytochrome P450 monooxygenases (CYP450), glutathione s-transferases (GSTs) and UDP-glucosyl transferase (GTs) in the resistant plants. Taken together, these data suggest that metabolic detoxification of fluroxypyr may be the mechanism of fluroxypyr resistance in Flur-R.

## 2. Materials and Methods

### 2.1 Plant Materials

In 2014, 171 kochia (*Bassia scoparia)* populations were collected from a field survey conducted in eastern Colorado (Westra et al. 2019). These populations were screened at a single doses of dicamba, fluroxypyr, and glyphosate to test for resistance. One population, Flur-R, was found to have a few individuals (<2%) surviving a single fluroxypyr dose of 157 g ae ha^-1^ (label rate for use in wheat) (Starane Ultra, Dow Agrosciences, Indianapolis, IN). Survivors at 157g ae ha^-1^ were selected and allowed to bulk pollinate. After two more generations of selection at both 157g ae ha^-1^ and 314 g ae ha^-1^, the surviving individuals were cross-pollinated. The progeny of these plants were found to be uniformly resistant to 314g ae ha^-1^ fluroxypyr. During the bulking stages, groups of three to four plants were planted in one gallon pots and covered with a pollination bags to allow for cross-pollination. Seed was harvested per pot, hand threshed and cleaned using an air-column blower. Seeds were stored at 4 C and planted in the spring in a greenhouse maintained at 25 C with a 16 h photoperiod. An inbred dicamba resistant/fluroxypyr susceptible population (9425) homozygous for a G127N mutation in the *IAA16* gene (LeClere et al. 2018; Preston et al. 2009) and a fluroxypyr susceptible field population from the 2014 eastern Colorado field study (J01-S) (Westra et al. 2019) were included in the dose response and single dose screening as susceptible controls.

### 2.2 Fluroxypyr and Dicamba Dose Response

Seeds of Flur-R, J01-S, and 9425 were planted in 1.5 cm^2^ 280-count plug flats. Plants were sub-irrigated and thinned down to one plant per cell. When plants were approximately 4-5 cm tall, uniform seedlings were transplanted to 4 cm^2^ plastic pots containing SunGro potting mix (American Clay Works Supply, Denver, CO). Plants kept in the greenhouse under conditions previously described. They were sub-irrigated once a week for three weeks until the plants reached 10 cm height. A randomized complete block design was used for each dose, with one plant per pot, four plants per dose and three replications for a total of 12 treated plants. The dose response for fluroxypyr included the following eight rates: 0, 20, 40, 80, 157, 314, 628, and 1,256 g ae ha^-1^ fluroxypyr (Starane Ultra, Corteva Agrisciences, Indianapolis, IN). The dicamba doses included 0, 35, 70, 140, 280 (1x), 560, 1120, and 2240 g ae ha^-1^ (Engenia, BASF, Research Triangle Park, NC) mixed with Induce (NIS, 0.25% v/v, Helena Agri-Enterprises, LLC, 24330 US-34 Greely, CO 80631). Applications were made with a DeVries Generation 4 Research Track Sprayer (DeVries Manufacturing, Hollandale, MN, 86956) equipped with a TeeJet (TeeJet Technologies, 1801 Business Park drive, Springfield, IL) 8002EVS nozzle calibrated to deliver 187 L ha^-1^. Plant height was measured by recording the distance in centimeters from the soil surface to the newest leaf in the apical meristem before treating and was measured again 30 days after treatment. Survival (dead or alive) was also recorded 30 days after treatment. An individual was considered “dead” if it displayed severe epinasty, stem thickening, yellowing, and had no new growth at the axillary or primary meristems after 30 days. An individual was considered “alive” if it displayed minimal to no epinasty or stem thickening, had no yellowing, and had new growth at the axillary or primary meristems after 30 days. Percent survival was chosen for fluroxypyr resistance assessment because while percent change in height can accurately differentiate between resistant and susceptible plants, for this population it did not accurately represent an actively growing plant in the individuals where axillary meristem growth was the primary source of regrowth.

For data analysis, the response variable “Percent Survival” was created by transforming binary data according to the equation:

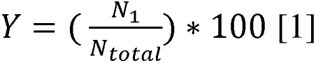

Where *Y* is the percent survival at each calculated dose, *N_1_* is the number of individuals marked as “alive” according to the parameters above. *N_total_* is the number of individuals per rate. The response variable “Percent Change in Height” over 30 days was normalized using the following equation:

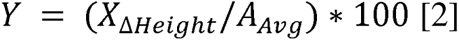

Where *Y* is the change in height as a percent of the 0 g ae ha^-1^ rate for each population. X_Δ*Height*_ is the change in height in centimeters for an individual from day 0 to 30 days, and *A_Avg_* is the average change in height for individuals at the 0 g ae ha^-1^ rate for the population being measured. The model used by the drm package in R did not converge for the J01-S or 9425 lines using “Percent Change in Height (% control)” as a response variable due to the non-sigmoidal behavior of the curve, so “Percent Survival” data were analyzed using a three-parameter log-logistic model (Knezevic et al. 2007), which was the best model by a lack-of-fit test from the drc package in R (R Core Team 2020) with the equation:

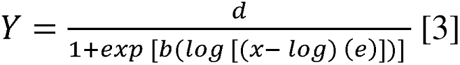

where *Y* is the percent survival 30 days after treatment, *d* is the upper limit parameter, *b* is the regression slope, *x* is the dose of either fluroxypyr or dicamba in g ae ha^-1^ and *e* is the dose at which 50% mortality is achieved (Table 1). The data were averaged per treatment and the standard error of the mean is presented per dose. “Rate” and “Population” were used as predictor variables and the experiment was repeated.

**Table 1.** Parameters for fluroxypyr dose-response data in kochia (*Bassia scoparia*) populations Flur-R, 9425, and J01-S. Parameters of the fluroxypyr and dicamba dose-responses for percent survival parameters are described in Equation 3 for Flur-R, fluroxypyr sensitive line J01-S and fluroxypyr sensitive/dicamba resistant line 9425. Flur-R shows a significant resistance factor ratio (R/S) of 36 and 40 relative to 9425 and J01, respectively. (*b,d*) Lower and upper limits of regression parameters, respectively. (*LD_50_*) The dose (g ae ha^-1^) of fluroxypyr where 50% mortality occurs for each population. (*R/S)* The ratio of resistant LD_50_ to either susceptible LD_50_ and associated p-values.

| Line | Fluroxypyr |  |  |  |  | Dicamba |  |  |  |  |
| --- | --- | --- | --- | --- | --- | --- | --- | --- | --- | --- |
|  | *b*<br>(± SE) | *d*<br>(± SE) | LD<sub>50</sub><br>(± SE) | R/S | P-value | *b*<br>(± SE) | *d*<br>(± SE) | LD<sub>50</sub><br>(± SE) | R/S | P-value |
|  | -----g ae ha<sup>-1</sup>----- |  |  |  |  | -----g ae ha<sup>-1</sup>----- |  |  |  |  |
| Flur-R | 7.3<br>(8.4) | 94.5 (3.6) | 720 (110.3) | 36-40 | <0.001 | 7.4 (5.2) | 100.0 (4.9) | 56 (8.5) | -- | -- |
| 9425 | 8.5<br>(37.7) | 100.0 (8.8) | 20<br>(1.5) | -- | -- | 85.1 (10.0) | 91.7 (2.72) | 415 (10.0) | 6-7 | <0.001 |
| J01-S | 3.1<br>(1.8) | 100.0 (8.8) | 18<br>(2.7) | -- | -- | 9.3 (8.5) | 100.0 (4.93) | 64 (5.4) | -- | -- |

### 2.3 Glyphosate, Atrazine, and Chlorsulfuron Single Rate Screening

Flur-R and J01-S seeds were planted in 4 cm^2^ plastic pots containing SunGro potting mix. Plants were sub-irrigated and thinned down to one plant per cell and kept at greenhouse conditions previously described. When plants were approximately 7 cm in height, plants were treated with one of the following herbicides (n=72 plants per herbicide): atrazine (Aatrex 4L, Syngenta, Greensboro, NC, 2240 g ai ha^-1^, 1% crop oil concentrate), chlorsulfuron (Telar XP, Bayer CropScience, St. Louis, MO, 137 g ai ha^-1^), or glyphosate (RoundUp Powermax, Monsanto Company, St. Louis, MO, 870 g ae ha^-1^, 2% w/v ammonium sulfate). All treatments were applied with the same equipment and nozzle type described above. Survival (dead or alive) was assessed 30 days after treatment. In a *post hoc* analysis, a random number generator was used to assign each of the 72 individuals to one of three blocks with n=24 to serve as replicates. Standard error of the mean was calculated using the standard deviation from this analysis.

### 2.4 Kompetitive allele specific PCR (KASP)

Approximately 200 mg of meristem tissue was harvested from 20 individuals each of Flur-R and 5 individuals for the mutant and wild type checks. Flur-R individuals were verified as resistant by spraying with 157 g ae ha^-1^ fluroxypyr. Tissue was put into a 1.5 mL Eppendorf tube and flash frozen in liquid nitrogen. DNA extraction protocol was adapted from Aboul-Maaty and Oraby (2019) using the established CTAB method. DNA purification was checked using a NanoDrop2000 and diluted to 5 ng uL^-1^. The FAM fluorophore (in bold) was added to the forward primer specific to the G127N *IAA16* double mutation (allele specific sequence in italics) endowing a protein change from wildtype GWPPV to NWPPV in kochia described by LeClere et al. (2018) (5’**GAAGGTGACCAAGTTCATGCT**TGTTCTTCAGGACACAAGTTGTA*AA*) and the HEX fluorophore (in bold) was added to the forward primer specific to the wild type sequence (in italics) (5’**GAAGGTCGGAGTCAACGGATT**TGTTCTTCAGGACACAAGTTGTA*GG*). One universal reverse primer (5’AGTTTGATCATCGGACGTCTTCTT) and the forward primers were designed with IDT PrimerQuest. The KASP protocol and specific mix ratios are published on protocols.io at dx.doi.org/10.17504/protocols.io.dm6gpj9njgzp/v1. Fluorescence was recorded at the end of every cycle. Fluorescence at the 35^th^ cycle was used for the allelic discrimination data. Data were plotted using GraphPad Prism version 8.4.2. Genotypes were assigned manually as homozygous wildtype, homozygous mutant, or heterozygous.

### 2.5 Plant Material for Fluroxypyr Absorption, Translocation, and Metabolism

Seeds from Flur-R and J01-S were sown into plug flats filled with SunGro potting mix and grown on a light shelf under 700 μmol m^-2^ s^-1^ of light at 25 C. When the plants reached 3-4 cm tall, 50 seedlings from each line were washed of soil in the roots and transplanted into a 25 mL Eppendorf tube filled with silica sand and fertilized with three granules of Osmocote. Uniform plants that were 4-5 cm tall and had recovered from transplanting were used in all subsequent absorption, translocation and metabolism experiments.

### 2.6 Fluroxypyr Absorption and Translocation

Flur-R (n=24) and J01-S (n=24) plants were sprayed with 157 g ae ha^-1^ fluroxypyr using a track sprayer as described in section 2.2. The third and fourth youngest leaves were protected from the broadcast application using aluminum foil. Immediately after applying fluroxypyr, the covered leaves of were treated with five 1 μL drops of the spray solution spiked with 3.1 kBq [^14^C]-fluroxypyr. Absorption and translocation were monitored over a 196 h time course, with time points at 6, 12, 24, 48, 96 and 192 hours after treatment (HAT). Four Flur-R and four J01-S were harvested at each timepoint. Treated leaves were removed and washed in 5 mL 90% water, 10% methanol, and 0.5% non-ionic surfactant. The leaf wash was mixed with 10 mL scintillation cocktail (Ecoscint XR) and radioactivity was measured using a liquid scintillation counter (LSC) (TRI-CARB 2300TR, Packard Instruments Co., USA). Roots were washed with 5 mL water. Root wash and the silica sand rinse solution were vortexed for 3 seconds, and 1 mL of the root wash and sand rinse mixture was added to 10 mL scintillation cocktail to measure root exudation. Plants were sectioned and separated as follows: above treated leaves, treated leaves, below treated leaves, and root biomass. Each separate plant part was dried and oxidized using a biological oxidizer (Model OX500, R. J. Harvey Instrument Co., USA). The released ^14^C-CO_2_ was collected by a ^14^C trapping cocktail (OX161, R.J. Harvey Instrument Co., USA). Radioactivity was quantified by LSC. One individual per timepoint was left intact, dried, and used for phosphor imaging (Typhoon Trio, GE Healthcare). Dried plants were separated and oxidized in the same manner as described above after imaging.

The experiment was repeated. Percent absorption and translocation was calculated as follows from Figueiredo et al. (2018) and maximum percent absorption was determined using a method described by Kniss et al. (2011) in R.

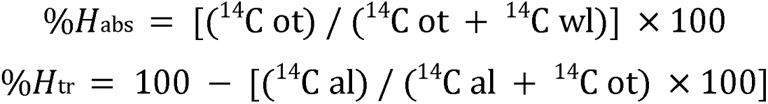

Where “%*H*_abs_” is percent absorption of [^14^C]-fluroxypyr ester, “^14^C ot” is the sum DPM from the oxidation of all plant parts and “^14^C ot + ^14^C wl” is the sum DPM from the oxidation of all plant parts and counts washed from the treated leaf. For herbicide translocation studies, “%*H*_tr_” is percent translocation of [^14^C]-fluroxypyr ester out of the treated leaf through the rest of the plant, “^14^C al” is the DPM [^14^C]-fluroxypyr ester counted in the treated leaf.

### 2.7 Fluroxypyr Metabolism

Fluroxypyr metabolism was evaluated by treating Flur-R and J01-S plants as perviouly described. Plants were sprayed with fluroxypyr while two leaves were protected from the spray solution. Those protected leaves were then treated with five 1 μL drops of the spray solution spiked with 25 kBq [^14^C]-fluroxypyr. The time course was the same as previously described with four repetitions per timepoint. Treated leaves were removed and washed as previously described. The washed treated leaves were placed back with the remaining whole-plant tissue and flash frozen in liquid nitrogen. Whole plants were then finely ground in a glass test tube with liquid nitrogen and a glass rod. Five mL extraction solution (90% water, 9% acetonitrile, 1% acetic acid) was added to each tube and samples were shaken for 30 min. The extraction solution was tranferred to a 0.45 µm filter tube which was rinsed with an additional 5 mL extraction buffer and centrifuged at ∼2600×g for 10 min to separate liquid from ground plant material. The extraction buffer that passed through the 0.45 µm filter was transferred to a C-18 cartridges preconditioned with 1 mL 100% acetonitrile (Waters Co., Sep-Pak Plus). Using a vacuum manifold, the extraction buffer was pulled throught the solid phase extraction cartridge. Bond solutes were eluted from the C18 cartridge with 5 mL 100% acetonitrile and samples were then evaporated to dryness in a fume hood. Solvent A (500 µL) consisting of 10% acetonitrile and 1% formic acid was added to each tube. Each sample was filtered through 25 µm nylon filters (Nalgene) into an injection vial with a 500 µL insert. High Pressure Liquid Chromatography (HPLC) (Hitachi Instruments, Inc.,) was used to separate radiolabelled fluroxypyr-ester, fluroxypyr acid and metabolites.. The HPLC was equipped with a C-18 column (4.6 mm by 150 mm column, Zorbax Eclipse XDB-C18, Agilent Technologies) and inline radio-detector (FlowStar LB 513, Berthold Technologies GmbH & Co.) with a YG-150-U5D solid scintillation flow cell (150 μL). The injection volume was 200 µL. Radiolabelled fluroxypyr-ester had a retention time of 9.8 min, while fluroxypyr acid had a retention time of 2.8 minutes (protocol published on protocols.io at dx.doi.org/10.17504/protocols.io.kqdg39yopg25/v1).

### 2.8 Plant Material and Treatment for RNA Sequencing

Seeds from lines Flur-R, 9425, and J01-S were treated as described above, in similar growing conditions. When the plants reached 7-10 cm tall, 20 of the most uniform seedlings from each line were treated as follows: All plants were sprayed with water and 0.01 g meristem tissue was harvested for the untreated RNA-sequencing timepoint. Tissue was flash frozen in liquid nitrogen. The same twenty plants per line were treated 24 h later with 157 g ae ha^-1^ fluroxypyr, the labeled rate to control kochia. Approximately 0.01 g of meristem tissue was harvested at 3 and 10 h after fluroxypyr treatment for the remaining two RNA-sequencing timepoints. Herbicide and water applications were made with a track sprayer as described in section 2.2. All plants were in the vegetative stage, except for one Flur-R individual and three J01-S individuals, which were in the early flowering stage at the time of tissue harvest. After 30 days, resistance response was measured and four individuals per timepoint per line were selected for RNA-sequencing.

### 2.9 RNA Extraction, Sequencing and Quantification

The RNA-sequencing experiment was conducted first by extracting total RNA following the protocols in the QIAGEN RNeasy plant mini kit in six batches containing two individuals of each line to minimize batch effects. The kit was used to extract RNA from the top three fully expanded apical meristem leaves of Flur-R, 9425, and J01-S of 5-7 cm tall kochia at 0, 3, and 10 h after 157 g ai ha^-1^ fluroxypyr treatment. Final elution volume was 30 μL. Total RNA samples were diluted to a range of 500-10,000 pg μL^-1^ for quality check using an Agilent ScreenTape.

Samples that had a RIN score above 6 were submitted to BGI Technologies for quality check following their sample submission guidelines. Following quality check by BGI, 30 samples were used for sequencing. From the total RNA, mRNA enrichment was performed by rRNA depletion. Reverse transcription of the mRNA was performed with random N6 primers followed by end repair and A-tail and adapter ligation to the fragments. After PCR amplification, single strand separation and single-strand circularization were conducted to sequence paired end 100 base pair fragments with the BGISEQ sequencing platform. In total, 2.8 billion reads were produced, resulting in 92-97 million 100 bp reads per sample.

### 2.10 Differential Expression and Variant Analysis

Individual fasta files were uploaded to the remote research computing resource SUMMIT (Jonathon Anderson 2017) and files were quality checked with FastQC (version 0.11.9). Adapters were trimmed by BGI Bioinformatics company after sequencing and quality check. Reads were aligned to the *Bassia scoparia* coding sequence version 2 (Hall et al. 2023) using HISAT2 (version 2.2.0 (Kim et al. 2019)). Reads were assigned to features using featureCounts in the Subread package (version 2.0.1, Liao et al. 2014). Differential expression was conducted with resultant reads for each gene feature using the DESeq2 package (version 1.28.1) in the statistical software R (version 4.0.2 (R Core Team 2020)). Reads were transformed to logarithmic fold change log2 and compared across biological replicates for each population. For each population, the untreated condition was compared to either the 3 or 10 h timepoint to determine expression. Mean normalized counts per gene, an adjusted pvalue of < 0.05, and log2 fold change > 0.5 were the pre-filtering parameters used by DESeq2 for optimal significant genes below the false discovery rate (FDR) of <0.05.

Sorted and indexed bam files were run through the variant calling software Platypus (version 0.8.1 (Rimmer et al. 2014)) to detect single and mono-nucleotide polymorphisms, short and long indels, as well as chromosome rearrangement. The output file was used with the software SnpEff (version 4.3 (Cingolani et al. 2012)) to annotate the variants called from Platypus and to provide effect predictions. Specific genes annotated with involvement in the auxin signaling pathway or metabolic herbicide resistance were targeted for variant analysis by checking chosen genes against a merged variant file for all individuals of each population. Presence or absence of variants were validated with Integrative Genomics Viewer (IGV) (Robinson et al. 2017).

## 3. Results

### 3.1 Fluroxypyr and Dicamba Dose Response

Flur-R was confirmed to be fluroxypyr resistant based on change in height (Figure 2A) and percent survival (Figure 2B) at 30 days after treatment (DAT), with 75% survival up to 628 g ae ha^-1^ of fluroxypyr (Figure 2B). Flur-R was approximately 40 times more resistant than the susceptible population J01-S and 36 times more resistant than 9425 (Table 1). The population 9425, which was previously reported to be fluroxypyr resistant (LeClere et al. 2018) was subsequently shown to have weak fluroxypyr resistance (Wu et al. 2020). Our results show 9425 had less than 25% survival at 157 g ae ha^-1^ fluroxypyr (Figure 2B) and had similar reduction in height as the known susceptible population J01-S (Figure 2A). The LD_50_ ratios for J01-S and 9425 were not statistically different from 1, indicating that 9425 is not resistant to fluroxypyr at field rates. Furthermore, Flur-R was susceptible to dicamba (Figure 2C), with 8% survival at 70 g ae ha^-1^ and an LD_50_ of 56 g ae ha^-1^. Flur-R is approximately seven times more susceptible than 9425 to dicamba (Table 1).

**Figure 2:**
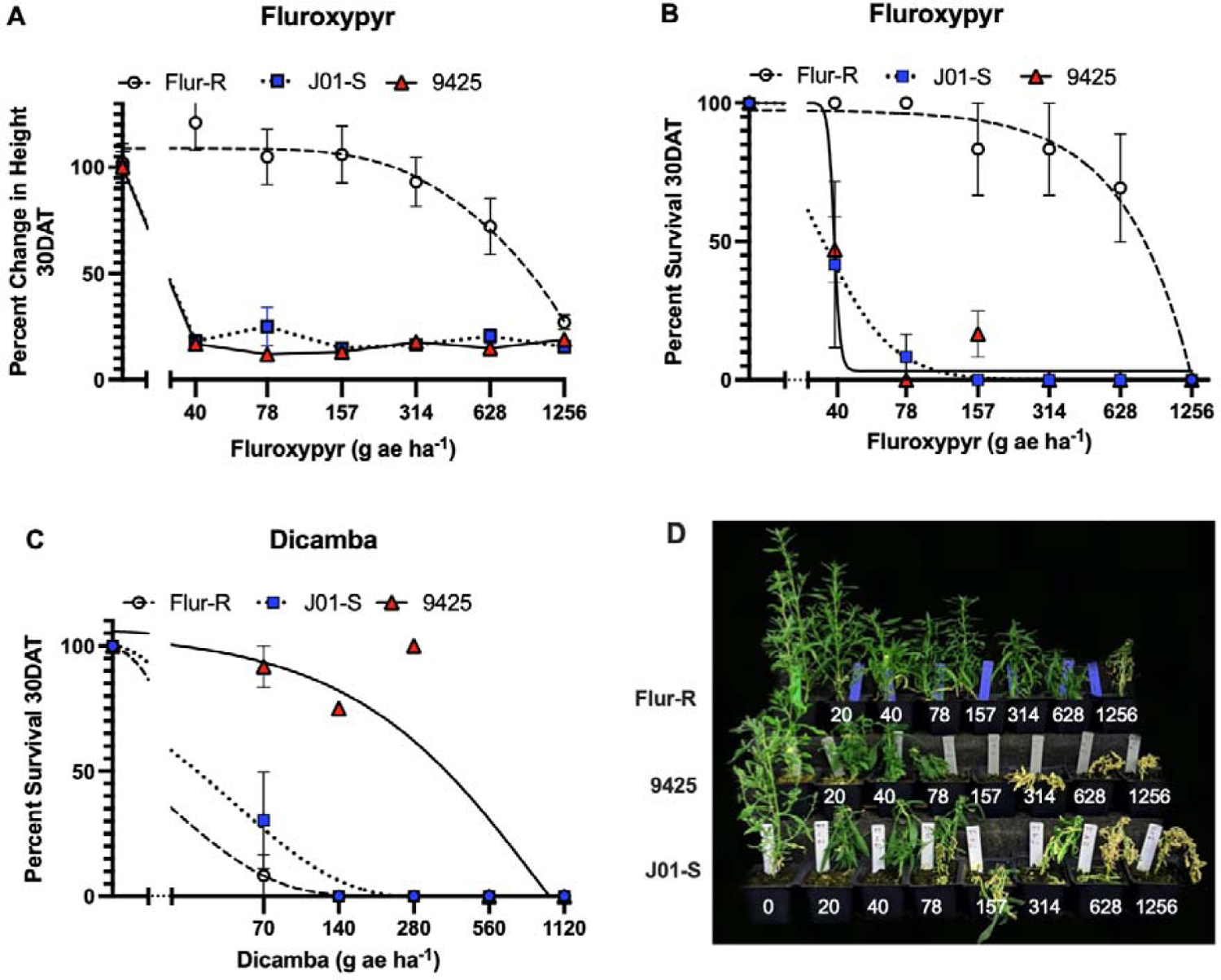
Dose response data for (A, B) fluroxypyr (no adjuvant) and (C) dicamba (+ 0.25% NIS) demonstrated fluroxypyr resistance and dicamba sensitivity in fluroxypyr resistant line Flur-R. X-axis is represented in a log10 scale. A. Percent change in height as a percent of the untreated control 30 days after treatment with fluroxypyr showed a 25% reduction in height in Flur-R at 628 g ae ha^-1^ (four times the label rate of 157 g ae ha^-1^). B. Percent survival for Flur-R with greater than 70% survival to fluroxypyr at 628 g ae ha^-1^ (LD_50_ = 720, p= <0.001). The population 9425 was susceptible to fluroxypyr (LD_50_ = 20 g ae ha^-1^, p <0.001). C. Flur-R was susceptible to dicamba (LD_50_ = 56 g ae ha^-1^, p <0.001) and the known dicamba-resistant line, 9425, was resistant to dicamba up to 280 g ae ha^-1^. Error bars represent SEM. D. Singular plants represent the average line response at each dose of fluroxypyr where 157 g ae ha^-1^ represents the label rate.

### 3.2 Glyphosate, Atrazine, and Chlorsulfuron Single Rate Screening

No Flur-R or J01-S individuals survived glyphosate (870 g ae ha^-1^) or atrazine (2240 g ai ha^-1^) treatments; however, 94% (± 0.5%) of the Flur-R population and 7% (± 0.5%) of J01-S individuals survived chlorsulfuron (137 g ai ha^-1^). This indicates that there is multiple resistance in this fluroxypyr resistant population. Two target site mutations were identified in a SNP analysis of RNA-sequencing data that confer ALS resistance, including a proline 197 to threonine mutation and a tryptophan 574 to leucine mutation in the *ALS* gene (Tranel and Wright 2002) (SI Figure 1).

### 3.3 KASP

Kompetitive allele specific PCR (KASP) was used to genotype individuals using allelic discrimination to determine whether or not Flur-R individuals contained the G127N *IAA16* mutation reported by LeClere et al. (2018). Specific fluorophore sequences were assigned to each forward primer, which generated a fluorescent signal to determine which allele was present in the kochia DNA sample. Relative Fluorescence Units (RFU) were measured to determine which of the fluorophore sequences amplified for each sample (Figure 3A). Of the twenty verified fluroxypyr-resistant individuals tested from the Flur-R population, 10 individuals displayed high RFU for the HEX labeled primer, indicating they had a homozygous wildtype genotype. There were six individuals that displayed approximately equal RFU for both alleles, indicating heterozygous individuals for G127N *IAA16*. Two known susceptible wild type controls were included (kochia lines J01-S, 7710), as well as one homozygous mutant resistant control (9425). These results indicate that the asparagine-127 *IAA16* mutant allele is not essential for fluroxypyr resistance, as most individuals were homozygous for the wildtype glycine-127 *IAA16* allele and were resistant to fluroxypyr. The dicamba resistance asparagine-127 *IAA16* allele is present and segregating in the Flur-R population.

**Figure 3:**
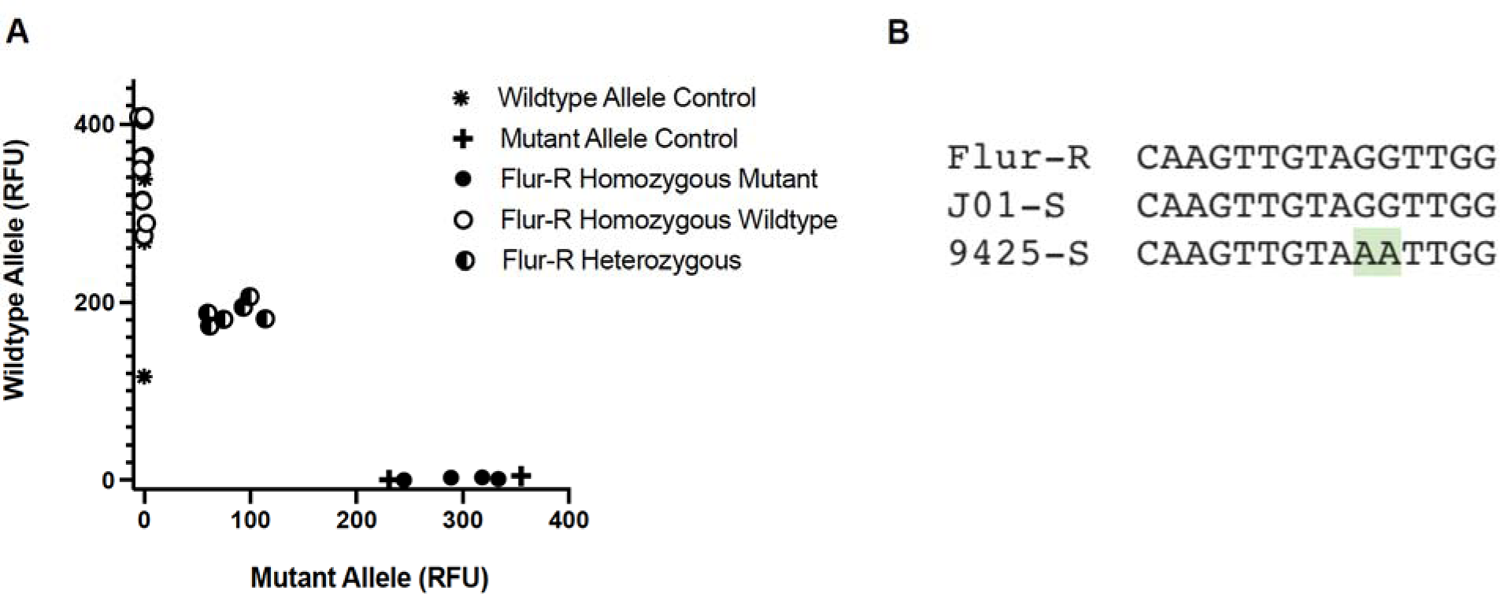
A. KASP assay with fluroxypyr resistant individuals sprayed with 157 g ai ha^-1^ fluroxypyr. Wildtype allele control lines were fluroxypyr and dicamba susceptible line 7710 and J01-S. Mutant allele control was homozygous dicamba resistant line 9425. B. Consensus sequenced based on KASP and previously published sequencing data at the mutation point of interest in J01-S, Flur-R and dicamba resistant line 9425 conferring resistance to dicamba as reported by LeClere et al. 2018. The mutation from GG to AA confers a G127N change.

### 3.4 Fluroxypyr Absorption, Translocation, and Metabolism

We investigated differences in [^14^C]-fluroxypyr ester absorption and translocation between fluroxypyr-resistant line Flur-R and fluroxypyr-susceptible J01-S. For each of the two lines, two meristem leaves per individual were treated with [^14^C]-fluroxypyr ester. Differential absorption and translocation were investigated by partitioning all individuals into four plant sections. Radioactivity was quantified in each section using biological oxidation and liquid scintillation counting. Maximum percent absorption of ∼3.33 kBq [^14^C]-fluroxypyr ester for Flur-R was 91.99% (±3.14), and for J01-S was 85% (±3.13). Percent recovery of radioactivity was >75% for all samples, except two samples per line at 192h which were ≥50%. No significant differences in maximum absorption between Flur-R and J01-S were found (pvalue = 0.155) (Figure 4A). The time (h) after treatment in which 90% of the herbicide is absorbed based on the model in R was not statistically different between 12 h (± 2.15) for Flur-R, and 9.7 h (± 2.34) for J01-S (pvalue = 0.47). There were no differences in translocation of [^14^C]-fluroxypyr ester from the treated leaf to the rest of the plant between Flur-R and J01-S (Figure 4B and C).

**Figure 4.**
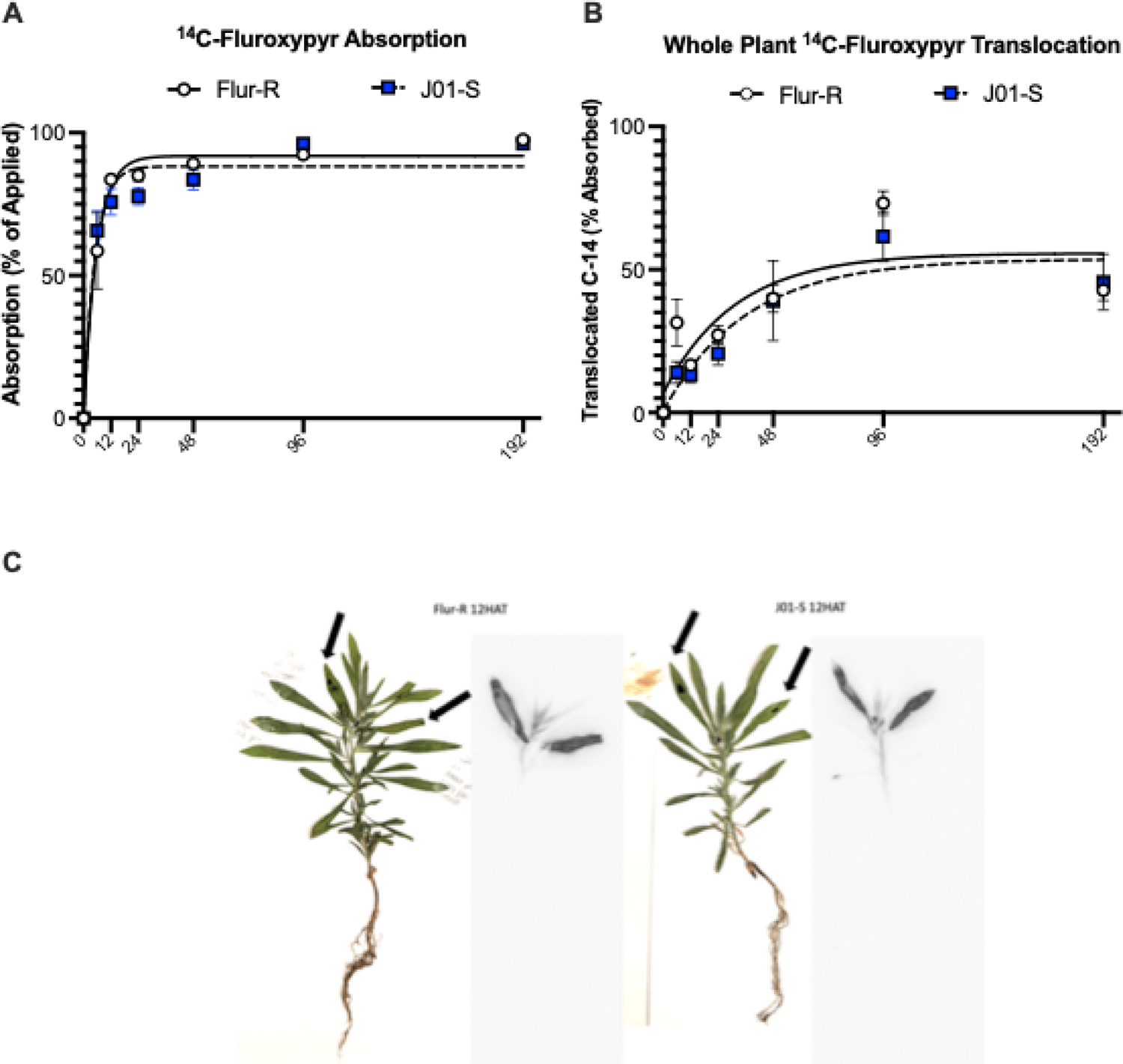
Whole plant absorption (A) and translocation (B) of fluroxypyr on resistant Flur-R line and susceptible line J01-S assessed over 6, 12, 24, 48, 96, and 192 h after treatment with [^14^C]-fluroxypyr ester. The absorption and translocation graphs depict mean percent absorption as percent of applied radiation and mean percent translocation as percent of absorbed radiation to account for slight variation in application rates, with error bars representing SEM. There were no differences in absorption or translocation of [^14^C]-fluroxypyr ester between Flur-R and J01-S. C. Pressed plant and phosphor-images showed translocation of [^14^C]-fluroxypyr ester in Flur-R (left) and J01-S (right) at 12 h, the time at which max absorption was at 90% in both lines. The black arrows mark the two treated meristem leaves on each individual. The phosphor image to the right of each pressed plant photo shows early-stage translocation of [^14^C]-fluroxypyr ester.

Whole plant metabolites were extracted for [^14^C]-fluroxypyr ester metabolism studies. Analysis of metabolites was conducted using an HPLC equipped with a C18 column. [^14^C]-fluroxypyr ester and acid standards were analyzed using an HPLC to determine retention time. Analysis of the proportion of [^14^C]-fluroxypyr ester in each population showed a significant difference at 12 h. The overall proportion of [^14^C]-fluroxypyr ester was lower in Flur-R than J01-S, supporting rapid conversion from the [^14^C]-fluroxypyr ester to biologically active [^14^C]-fluroxypyr acid or other [^14^C]-fluroxypyr metabolites (Figure 5). In Flur-R, the high amount of [^14^C]-fluroxypyr acid at 12 h was significantly reduced by 48 h, showing rapid conversion from to other fluroxypyr metabolites (Figure 5B). At 96 h and 192 h, the proportion of unknown metabolites numbered 4 and 2 were higher or uniquely present in Flur-R compared to J01-S (Figure 5D and F). This suggests that formation and flux of these metabolites is catalyzed by a process that is more active in Flur-R than J01-S and may play a role in reducing concentrations of phytotoxic [^14^C]-fluroxypyr acid.

**Figure 5.**
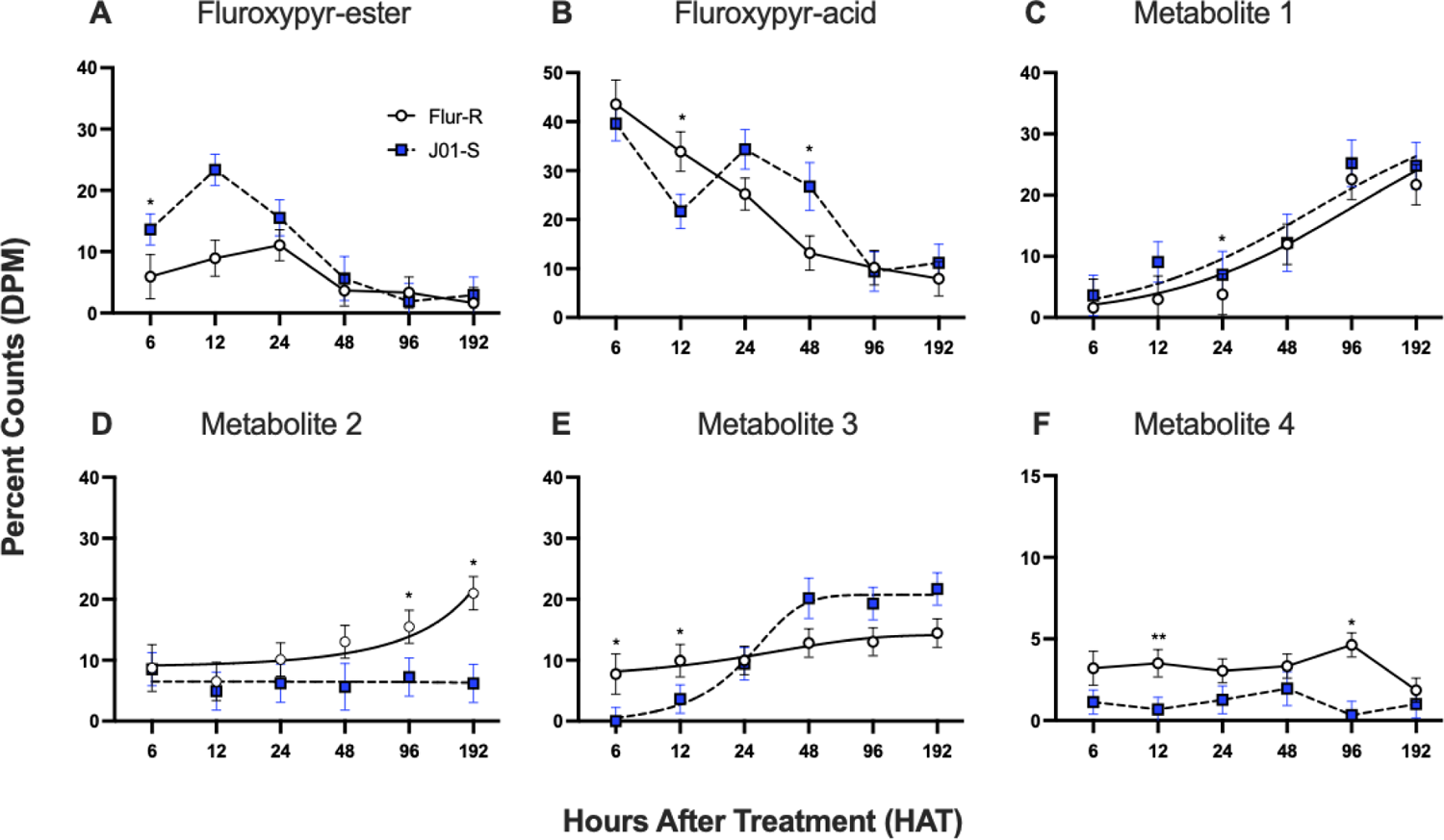
HPLC fluroxypyr parent and metabolite profiles over a 192 h time-course in fluroxypyr resistant kochia (*Bassia scoparia)* line Flur-R and fluroxypyr susceptible J01-S. A. Parent compound, [^14^C]-fluroxypyr ester (9.5 min retention). B. Biologically active compound, [^14^C]-fluroxypyr acid (7.7 min retention). C. Unknown metabolite 1 (4.5 min retention). D. Unknown metabolite 2 (5.8 min retention). E. Unknown metabolite 3 (6.4 min retention). F. Unknown metabolite 4 (7.2 min retention). * P<0.05, ** P≤0.005, error bars represent SEM.

### 3.5 Differential Expression Analysis

To analyze the transcriptome of Flur-R, we sequenced RNA from 4 plants each of fluroxypyr resistant Flur-R, and two fluroxypyr susceptible lines J01-S and 9425. BGI Seq was used to obtain between 91 and 95 million clean reads per sample (BGI Bioinformatics, San Jose, CA). Q20 scores were between 96–98%. Alignment was made to the coding sequence of the *Bassia scoparia* genome assembly version 2 (Hall et al. 2023) using HISAT2 (version 2.1.0), and alignment ranged between 59 - 63% for all individuals. Percent unmapped reads ranged between 46 – 51%, and percent uniquely mapped genes ranged from 43 – 48%. Approximately 4% of reads were multi-mapped (SI Table 1). Following alignment and differential expression with DESeq2, a Wald test was used to obtain p-values, which were adjusted using the Benjamini-Hochberg method. Filtering parameters included samples with an adjusted p-value < 0.05 and log2 fold change > 0.5. The false discovery rate (FDR) was < 0.05. We identified 231 unique genes that had higher expression in Flur-R compared to both 9425 and J01-S at the untreated timepoint (Figure 6). Because we identified differential metabolism in Flur-R, we explored the hypothesis that genes related to herbicide metabolism may have differential regulation or be highly expressed at the untreated timepoint in Flur-R. Of these 231 highly expressed genes in Flur-R at the untreated timepoint, there were six ABC transporters of both class B and G, including genes homologous to *ABCG31-like* (*Bs.00g217020.m01*), two similar ABCB28 annotated genes (*Bs.00g454440.m01, Bs.00g282300.m01*), two isoforms of *ABCG34* (*Bs.00g184080.m01, Bs.00g184080.m02*), and *ABCG29* (*Bs.00g251290.m01*). There were five CYP450 annotated genes between two families, the CYP71 family (*CYP82D47* [*Bs.00g486870.m01*], *CYP96A15* [*Bs.00g541440.m01*], *CYP71D10/11* [*Bs.00g051830.m01*], Ent-kaurene oxidase [*Bs.00g184110.m01*],) and the CYP85 family (*CYP90C1/D1* [*Bs.00g245700.m01*]). Several types of glucosyltransferases were expressed, such as UDP-glucosyltransferase 73B2 (*Bs.00g142060.m01*), two isoforms of UDP-glucuronosyl/UDP-glucosyltransferase 89A2-like (*Bs.00g480980.m01, Bs.00g480980.m02*), and UDP-glycosyltransferase 87A1 (*Bs.00g061050.m01*) (Table 2).

**Figure 6.**
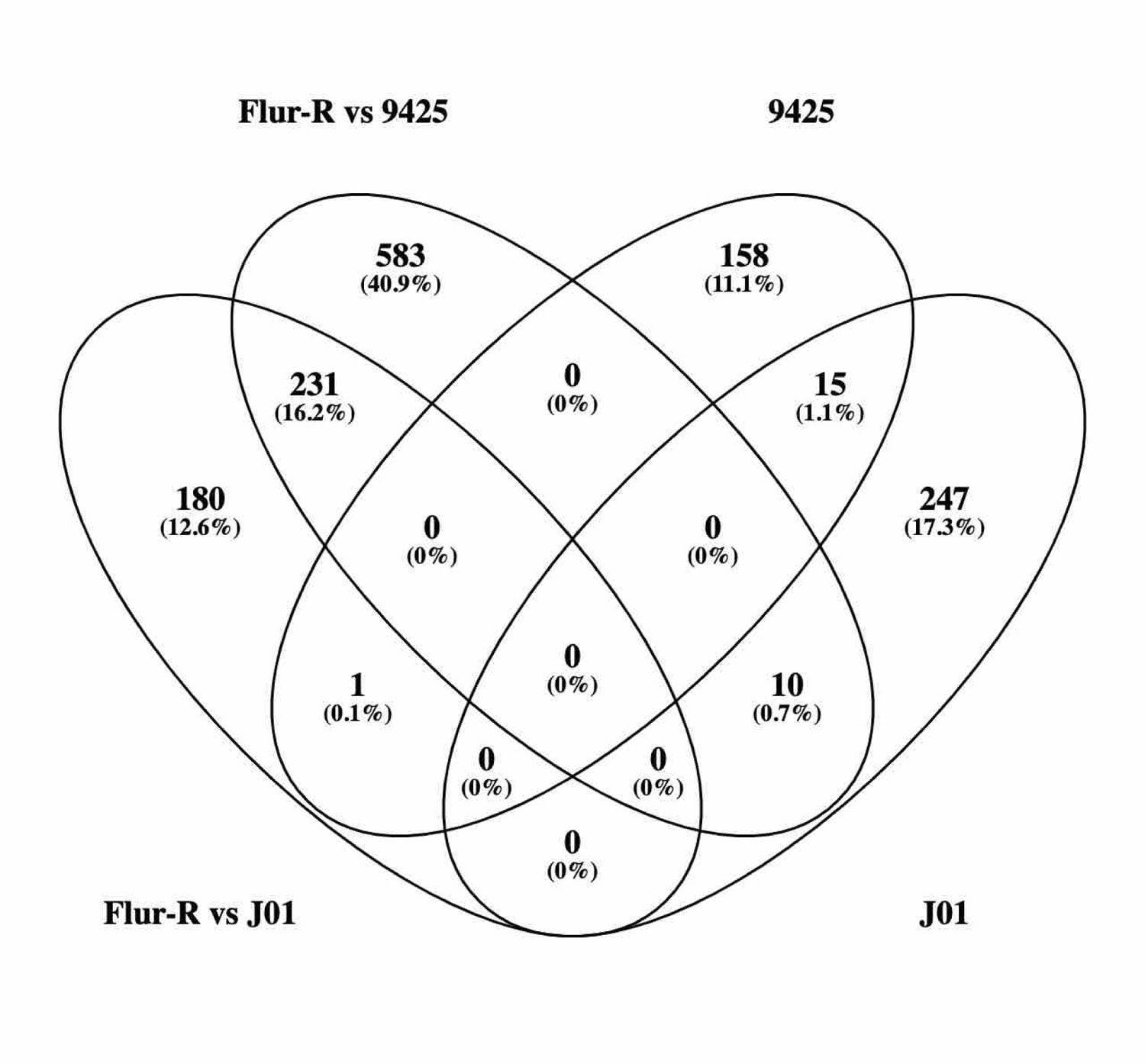
Venn diagram of upregulated genes between the untreated condition in Flur-R compared to both untreated conditions in fluroxypyr susceptible lines 9425-S and J01-S in DESeq2 (Flur-R vs 9425; Flur-R vs J01). Genes upregulated in both 9425-S and J01-S compared to Flur-R in DESeq2 are represented by their singular line name in the diagram (J01, 9425). Overlapping ovals represent genes that are commonly expressed at the untreated condition between comparisons.

**Table 2.** Genes with higher expression at the untreated timepoint in kochia (*Bassia scoparia*) line Flur-R compared to 9425 and J01-S lines at the untreated timepoint. Raw normalized counts and Log2 fold change for highly expressed ABC transporters, UDP glucosyltransferases, and cytochrome P450 monooxygenases in the fluroxypyr-resistant population Flur-R compared to either susceptible population 9425 or J01-S. Genes which are higher expressed in Flur-R compared to both susceptible populations and are denoted with † and represented with the normalized count and fold change comparison to 9425. Log2 Fold Change was calculated in DESeq2, log2 fold change standard error and adjusted *p*-value were also calculated in DESeq2. The Wald-test obtained *p*-values were adjusted using the Benjamini-Hochberg method. The FDR was < 0.05.

| Gene ID | Mean of normalized counts |  | Fold Change | Log2 Fold Change (±SE) | P-value (adjusted) | Gene Description |
| --- | --- | --- | --- | --- | --- | --- |
|  | Flur-R | 9425 |  |  |  |  |
| Bs.00g184080.m02 | 2258 | 3 | 753 | 9.70 (0.75) | 1.85E-32 | ABC-G 34 Isoform 2 |
| Bs.00g184080.m01 | 2147 | 3 | 716 | 9.63 (0.75) | 9.86E-32 | ABC-G 34 Isoform 1 |
| Bs.00g217020.m01† | 63 | 0 | 63 | 8.98 (2.85) | 1.31E-06 | ABC-G 31-like |
| Bs.00g142060.m01 | 38 | 0 | 38 | 8.02 (2.81) | 6.52E-05 | UDP-glucosyltransferase 73B2 related |
| Bs.00g184110.m01 | 133 | 2 | 67 | 6.24 (0.96) | 3.59E-08 | CYP701 subfamily (Ent- Kaurene Oxidase) |
| Bs.00g282300.m01† | 108 | 6 | 18 | 5.18 (1.47) | 0.0028 | ABC-G 28-like |
| Bs.00g480980.m01† | 510 | 14 | 36 | 5.01 (0.63) | 2.11E-11 | UDP-glucuronosyl/UDP-glucosyltransferase 89A2-like |
| Bs.00g480980.m03† | 538 | 15 | 36 | 4.99 (0.63) | 2.72E-11 | UDP-glucuronosyl/UDP-glucosyltransferase 89A2-like |
| Bs.00g541440.m01 | 232 | 10 | 23 | 4.21 (0.72) | 2.00E-06 | CYP96A15 |
| Bs.00g051830.m01 | 233 | 5 | 47 | 4.03 (1.48) | 0.0231 | CYP71D10/11 |
| Bs.00g454440.m01† | 276 | 52 | 5 | 2.28 (0.44) | 0.0003 | Putative ABC-B 28-like |
| Bs.00g061050.m01 | 6018 | 1187 | 5 | 2.08 (0.59) | 0.027 | UDP-glycosyltransferase 87A1 related |
| Bs.00g251290.m01 | 1462 | 341 | 4 | 1.98 (0.38) | 0.0008 | ABC-G 29-like |
| Bs.00g245700.m01 | 417 | 114 | 4 | 1.72 (0.40) | 0.0113 | CYP90C1/D1 (3-Epi-6-Deoxocathasterone 23-Monooxygenase) |
|  | Flur-R | J01-S |  |  |  |  |
| Bs.00g486870.m01 | 1847 | 61 | 31 | 4.85 (0.55) | 1.29E-13 | CYP82D47-like |
| Bs.00g142720.m01 | 8246 | 3606 | 2 | 1.17 (0.15) | 0.0001 | 7-deoxyloganetin glucosyltransferase-like 85A23 |

When analyzing the treated timepoints within Flur-R, J01-S, and 9425, 188 (3 h after treatment [HAT] vs untreated) and 300 genes (10 HAT vs untreated) were upregulated in response to fluroxypyr treatment in all three lines (Figure 7A and B). Of those shared upregulated genes, auxin responsive genes encoding proteins such as GH3.2 (*Bs.00g477580.m01*), Ethylene responsive transcription factors, Small auxin-up RNAs (SAURs), Aux/IAAs, and ACS (*Bs.00g478760.m01*) were among them, indicating that all three kochia lines perceived fluroxypyr and had transcriptional activation of these auxin responsive genes following fluroxypyr treatment (SI Figure 2). The Ethylene responsive transcription factors, *GH3*, and *ACS* were in the top 20 genes with the highest fold change through the 3 HAT vs untreated and 10 HAT vs untreated timepoints in Flur-R, J01-S, and 9425 (Tables 3, 4, 5). Two isoforms of the IAA cellular transporter PIN were upregulated in 9425 at 10 HAT (*Bs.00g190770.m01* and *Bs.00g190770.m02*), but the response in Flur-R and J01-S did not meet the differential expression filtering criteria and therefore the response was not statistically different following fluroxypyr treatment (Figure 8). Within the Flur-R line at 3 HAT vs untreated, there were 278 uniquely upregulated genes and 303 at 10 HAT vs untreated (Figure 7A and B). Some unique auxin-induced genes such as *SAURs* and *ARF11* were upregulated in Flur-R, but six additional ABC transporters of class G, two ABC transporters of class C, one ABC transporter of class A, six additional UDP-glucosyltransferases (GTs), and three sugar transporters were upregulated following fluroxypyr treatment (expression data not shown). CYP450s *CYP81B2* (*Bs.00g431990.m01*), *CYP82D47*, and *CYP71A9* (*Bs.00g241110.m01*) were induced by fluroxypyr treatment, as well as four Glutathione s-transferases (GSTs) in Flur-R at 3 and 10 HAT compared to the untreated timepoint.

**Figure 7.**
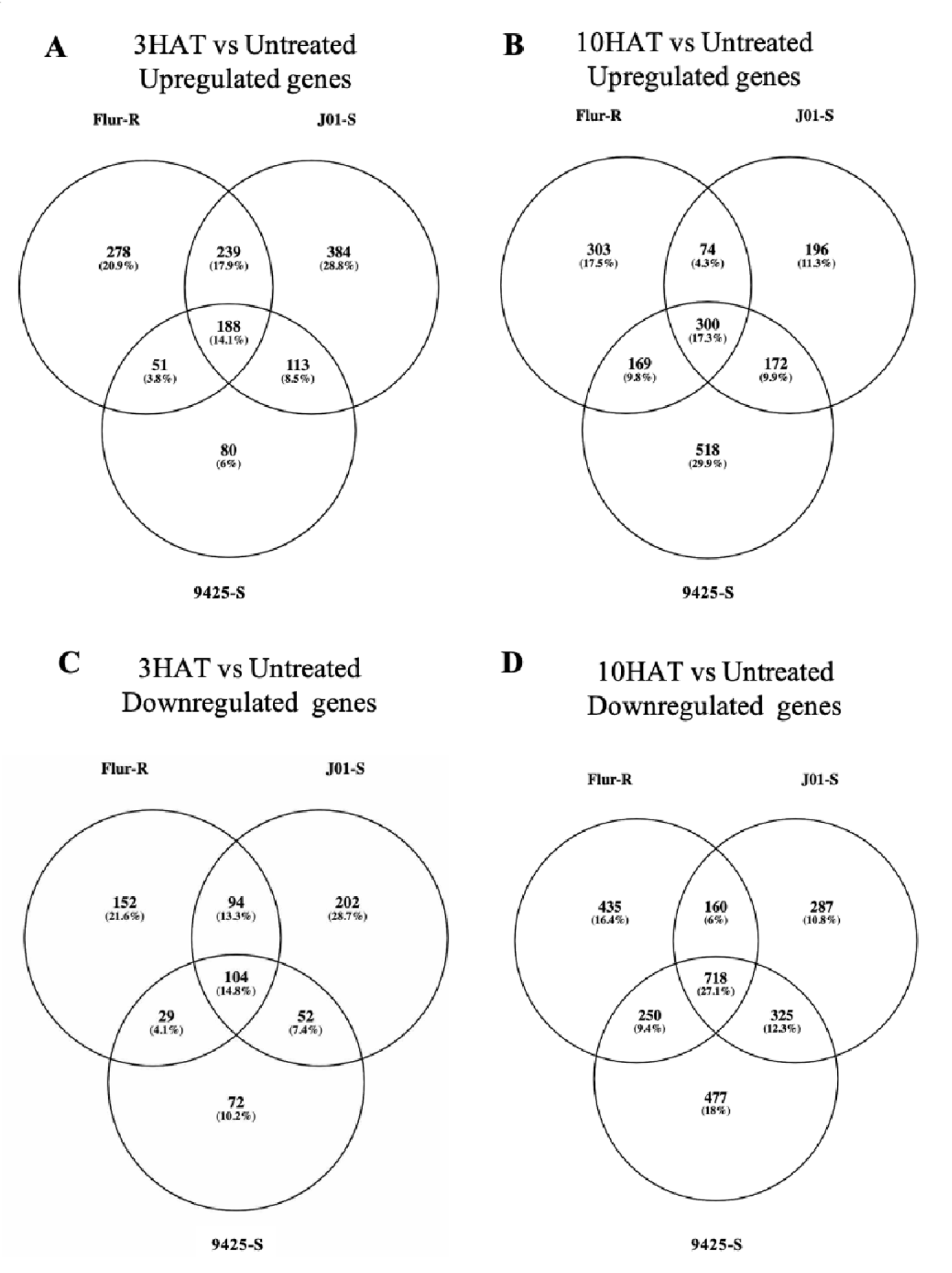
Number of transcripts that were either up or down regulated between the untreated condition and either 3 or 10 h after treatment (HAT) with fluroxypyr in fluroxypyr-resistant line Flur-R and susceptible lines 9425 and J01-S. A. Shared and uniquely upregulated genes at 3 HAT among and between all three lines. B. Shared and uniquely upregulated genes at 10 HAT among and between all three lines. C. Shared and uniquely downregulated genes at 3 HAT among and between all three lines. D. Shared and uniquely downregulated genes at 10 HAT among and between all three lines.

**Figure 8.**
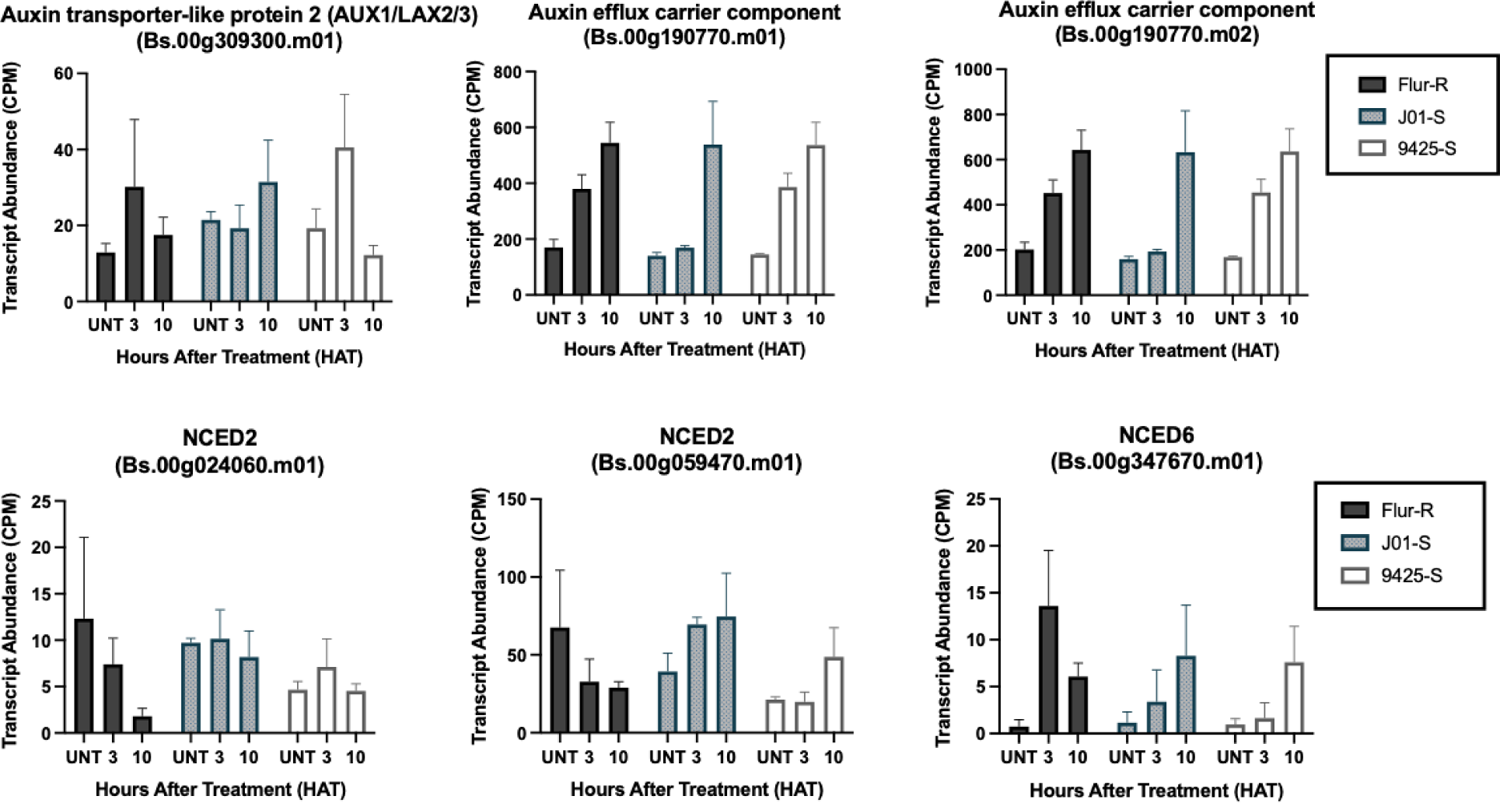
Expression profiles for auxin induced influx and efflux transporters and NCED in fluroxypyr-resistant kochia (*Bassia scoparia*) Flur-R, susceptible J01-S, and susceptible 9425 following differential expression analysis of RNA-Seq data. X-axis treatments: untreated, 3 h after treatment (HAT), and 10 HAT grouped by kochia line. Normalized counts on the y-axis were a result of the DESeq2 function and model fitting in R package “DESeq2”. Both isoforms of the auxin efflux carrier component were upregulated in response to fluroxypyr in the 9425 line. There were no differences in expression for the Aux/LAX transporter. NCED6 was induced at both 3 HAT and 10 HAT in Flur-R, while NCED2 was downregulated in Flur-R.

**Table 3.**
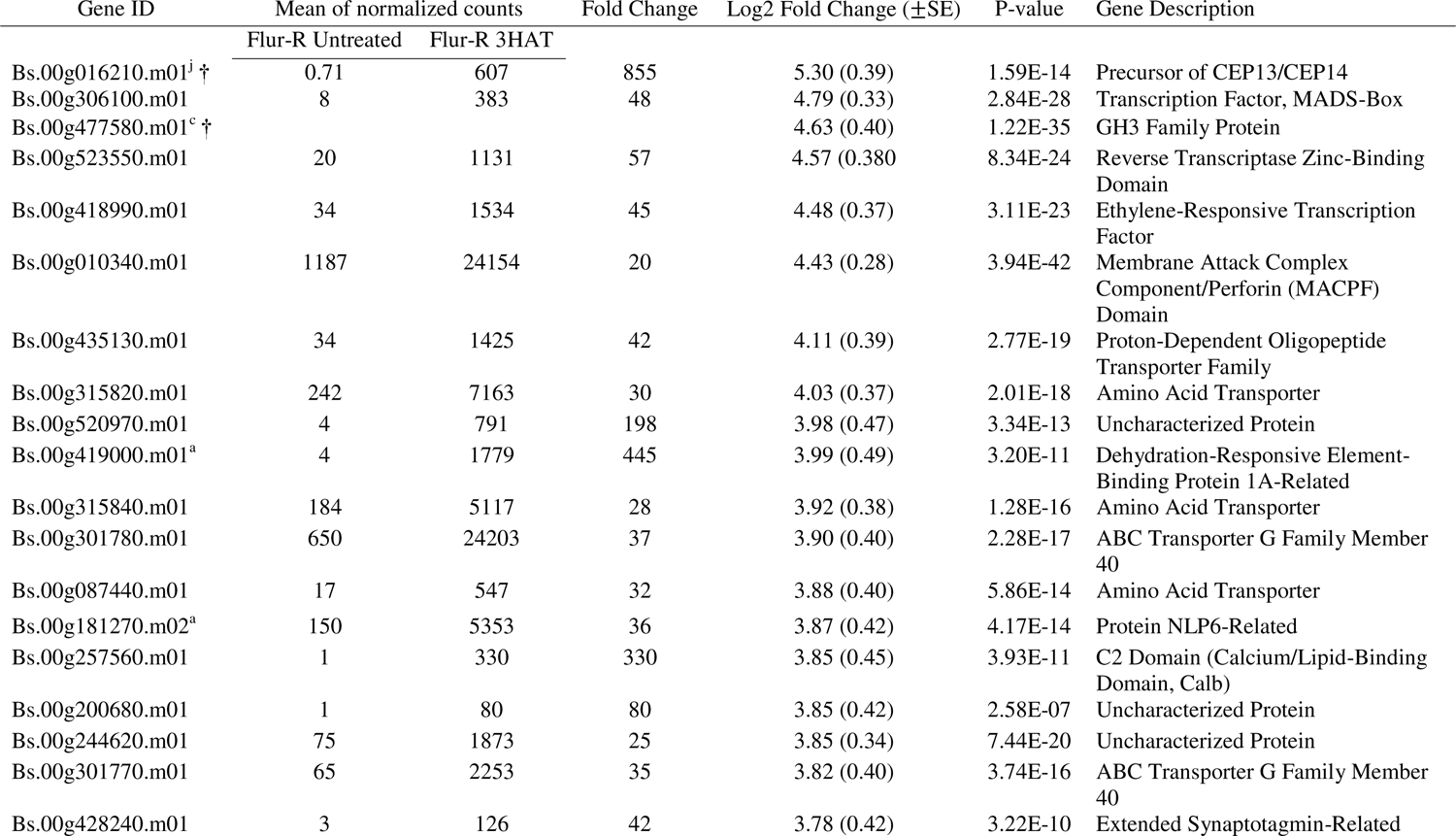
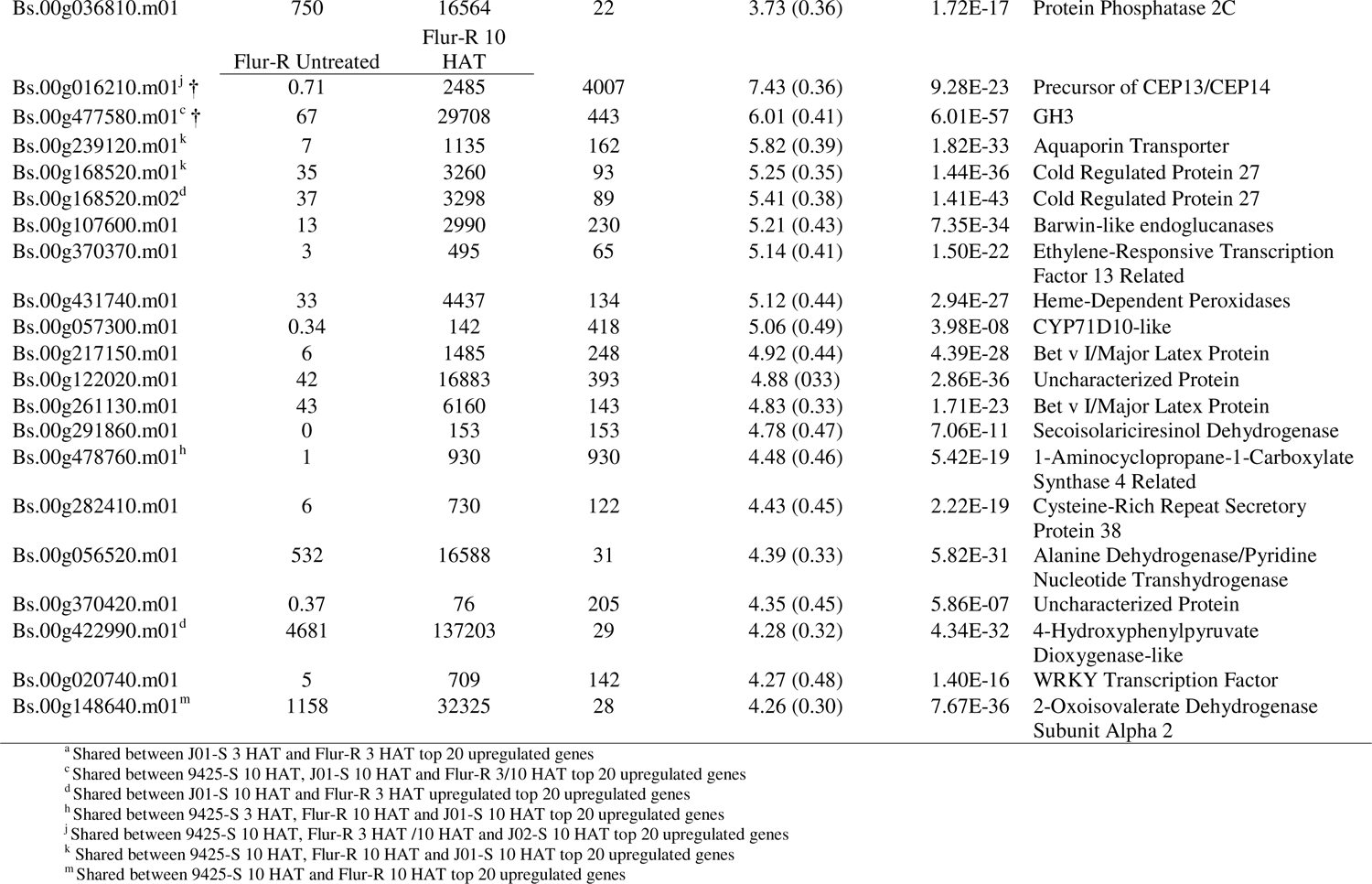
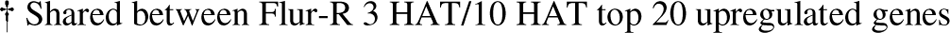
Top 20 upregulated genes in fluroxypyr-resistant kochia (*Bassia scoparia*) line Flur-R at 3 h after treatment (HAT) and 10 HAT compared to the untreated timepoint. Fold change was calculated using the mean of normalized counts, which was produced using the DESeq2 package in R. Log2 Fold Change was calculated in DESeq2, log2 fold change standard error and adjusted *p*-value were also calculated in DESeq2. The Wald-test obtained *p*-values were adjusted using the Benjamini-Hochberg method. The FDR was < 0.05.

**Table 4.**
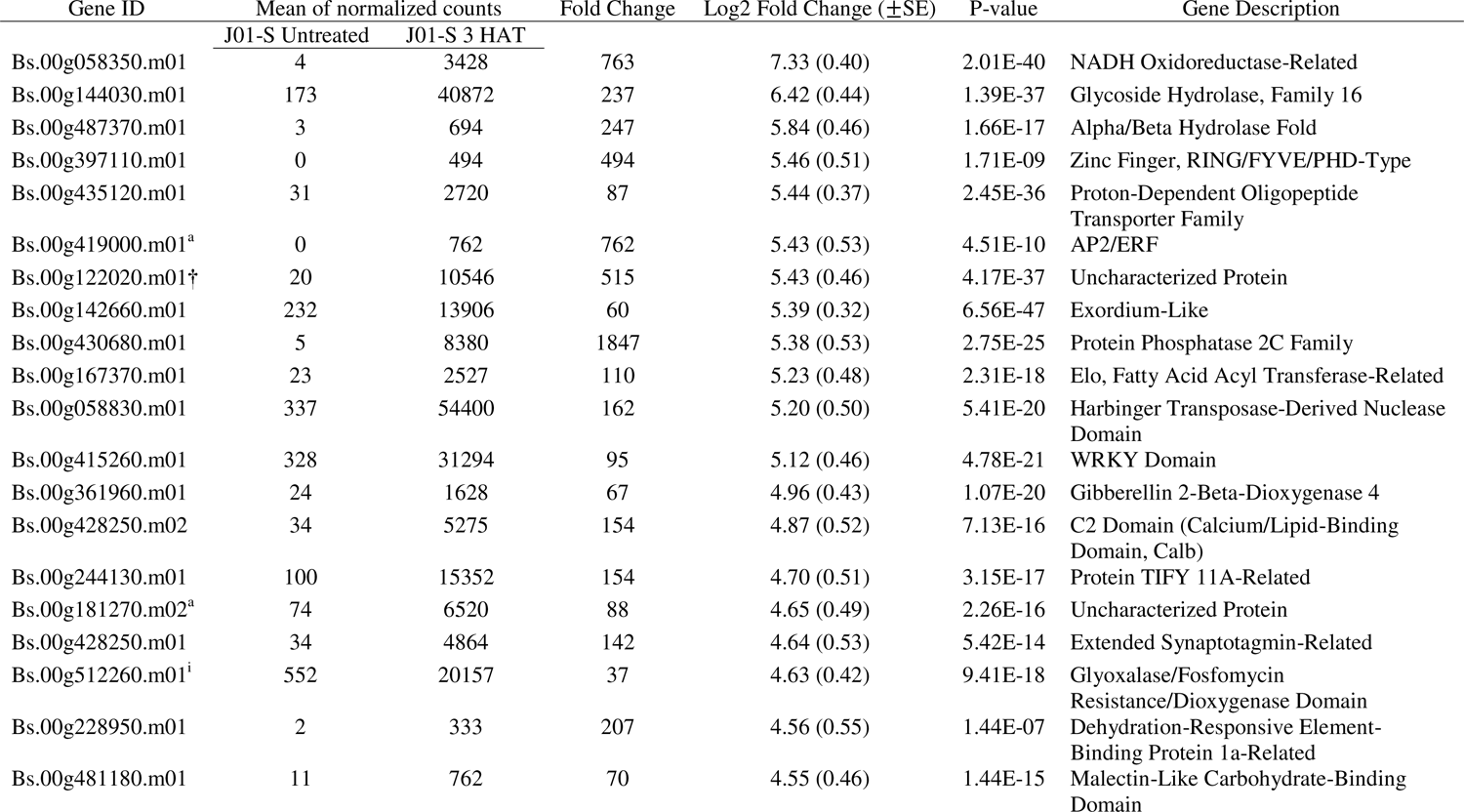
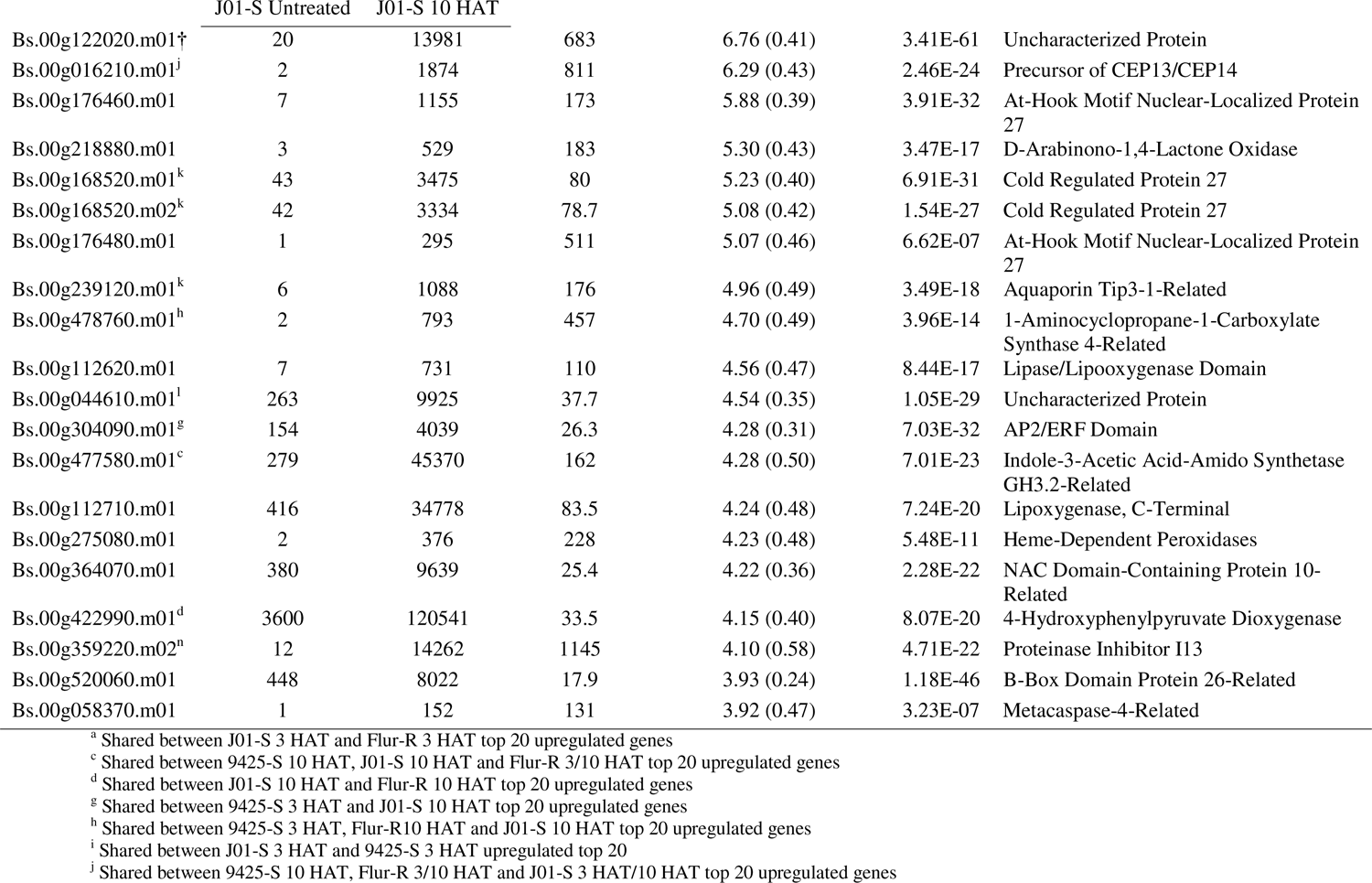
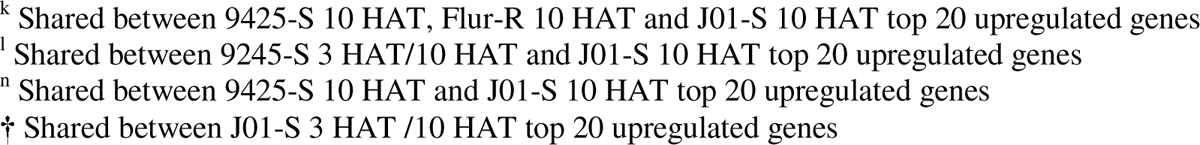
Top 20 upregulated genes in fluroxypyr-susceptible kochia (*Bassia scoparia*) line J01-S at 3 h after treatment (HAT) and 10 HAT compared to the untreated timepoint. Fold change was calculated using the mean of normalized counts which was produced using the DESeq2 package in R. Log2 Fold Change was calculated in DESeq2, log2 fold change standard error and adjusted *p*-value were also calculated in DESeq2. The Wald-test obtained *p*-values were adjusted using the Benjamini-Hochberg method. The FDR was <0.05.

When the downregulated 3 and 10 HAT timepoints were contrasted with the untreated timepoint within each line, Flur-R, J01-S, and 9425 identified 104 and 718 common fluroxypyr downregulated genes for the 3 and 10 HAT vs untreated timepoints, respectively (Figure 7C and D). Twelve of these shared genes were related to photosystem I and II at 10 HAT. Transcripts encoding key proteins related to photosynthetic electron transport such as Chlorophyll A-B binding protein (*Bs.00g240870.m01, Bs.00g240870.m02*) and ATP synthase (*Bs.00g432500.m01*) were downregulated in all three lines and were present in the top 20 downregulated genes for all three lines (Tables 6, 7, 8). Two chlorophyll biosynthesis regulator genes encoding Early light induced protein-1 (*Bs.00g421070.m01, Bs.00g420960.m01*) were uniquely downregulated and among the genes with the highest downregulation in both susceptible lines. These proteins play a role in preventing oxidative stress and excess accumulation of free chlorophyll (Hutin et al. 2003). Additionally, four Cellulose synthase genes (*Bs.00g015170.m01, Bs.00g015170.m02, Bs.00g056700.m01, Bs.00g260720.m01*) were downregulated in both susceptible lines and are of interest due to the role cellulose plays in cell wall structural support. Genes uniquely downregulated at 10 HAT in Flur-R included two additional photosystem II subunit genes (*Bs.00g059220.m01, Bs.00g338570.m01*) as well as genes encoding several synthases such as Terpene synthase (*Bs.00g074880.m01*), Polyprenyl synthetase (*Bs.00g449610.m01*), Strictosidine synthase (*Bs.00g057800.m01*), Phosphomethylpyrimidine synthase (*Bs.00g253210.m01*), Aminodeoxychorismate (ADC) synthase (*Bs.00g135570.m01*), and ABA biosynthesis gene *NCED2* (*Bs.00g024060.m01*).

**Table 5.**
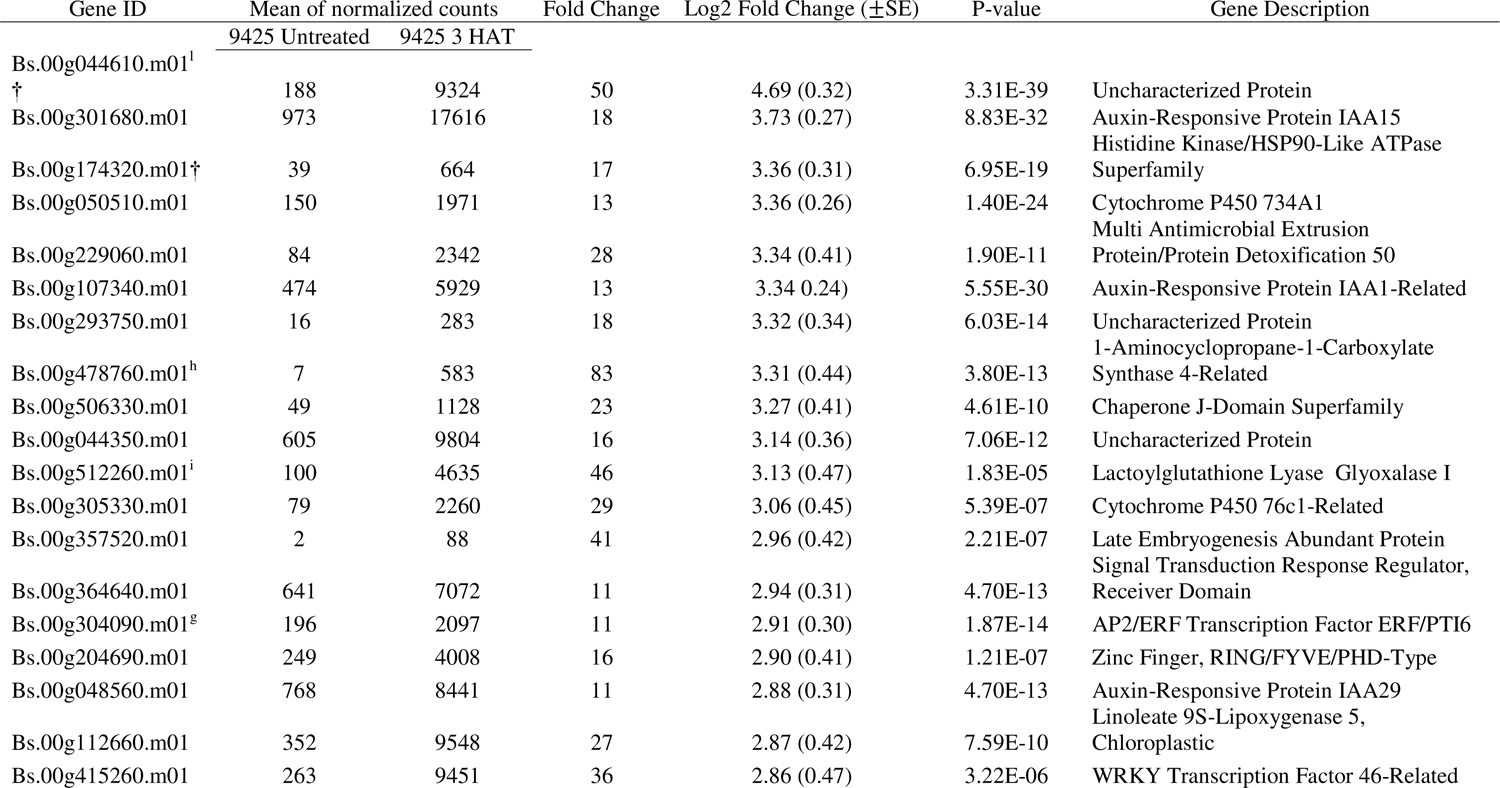
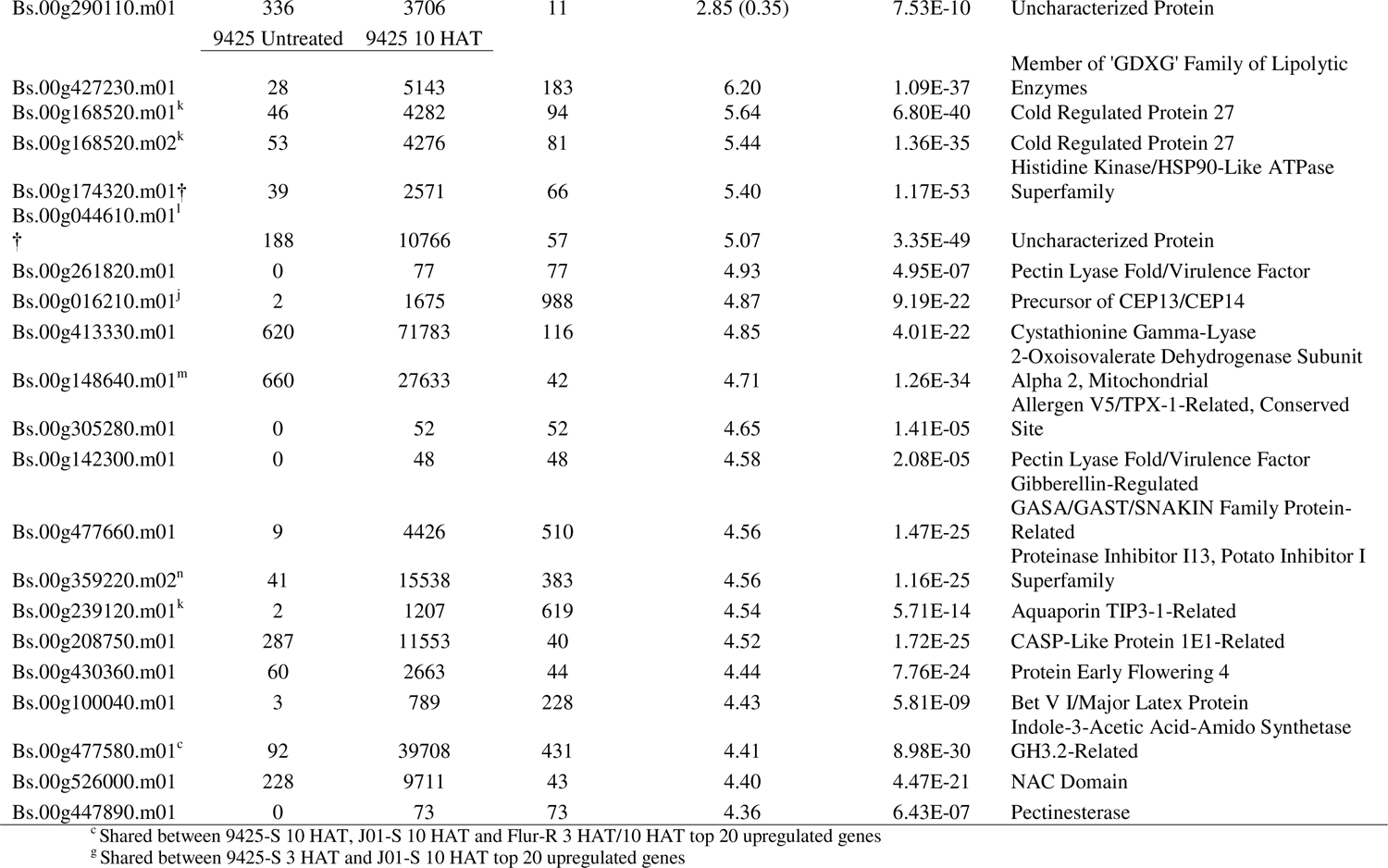
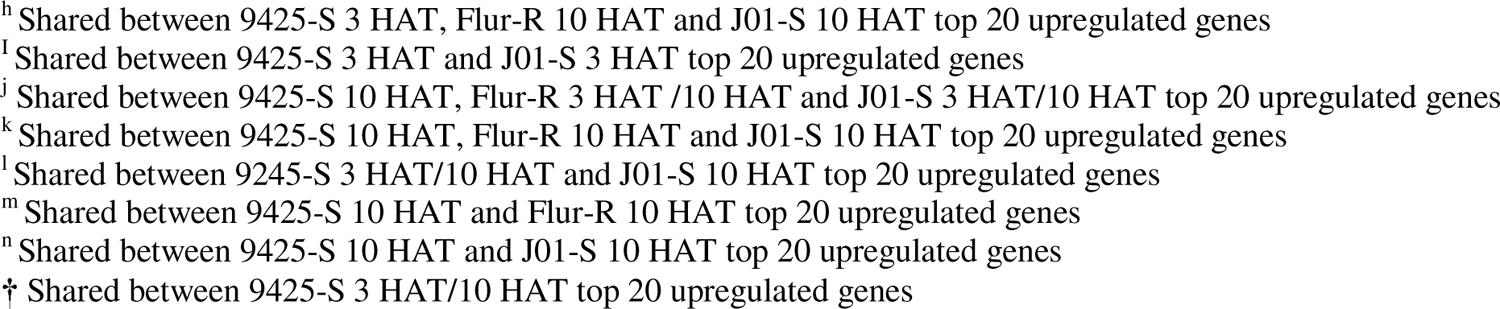
Top 20 upregulated genes in fluroxypyr-susceptible kochia (*Bassia scoparia*) line 9425 at 3 h after treatment (HAT) and 10 HAT compared to the untreated timepoint. Fold change was calculated using the mean of normalized counts which was produced using the DESeq2 package in R. Log2 Fold Change was calculated in DESeq2, log2 fold change standard error and adjusted *p*-value were also calculated in DESeq2. The Wald-test obtained *p*-values were adjusted using the Benjamini-Hochberg method. The FDR was <0.05.

**Table 6.**
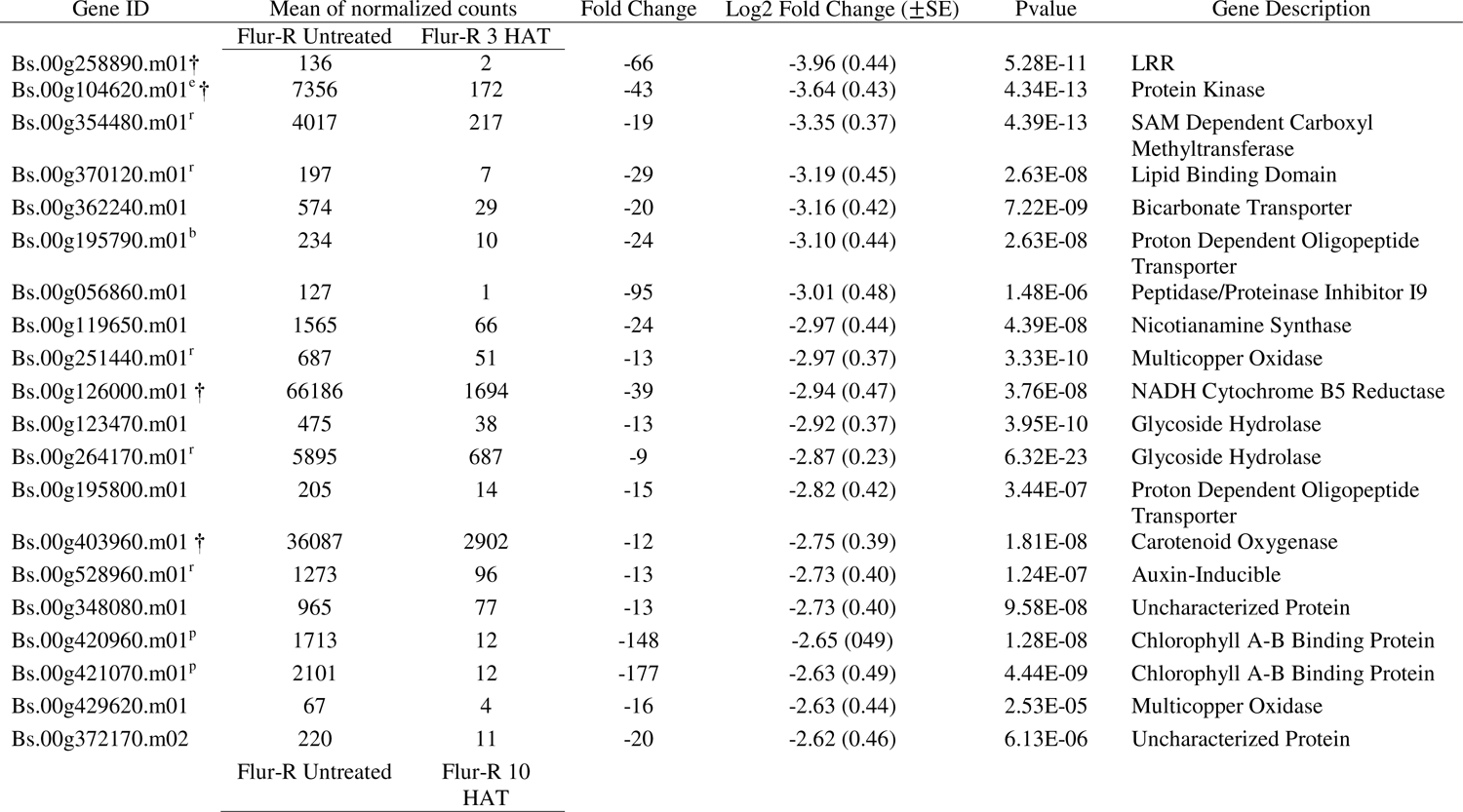
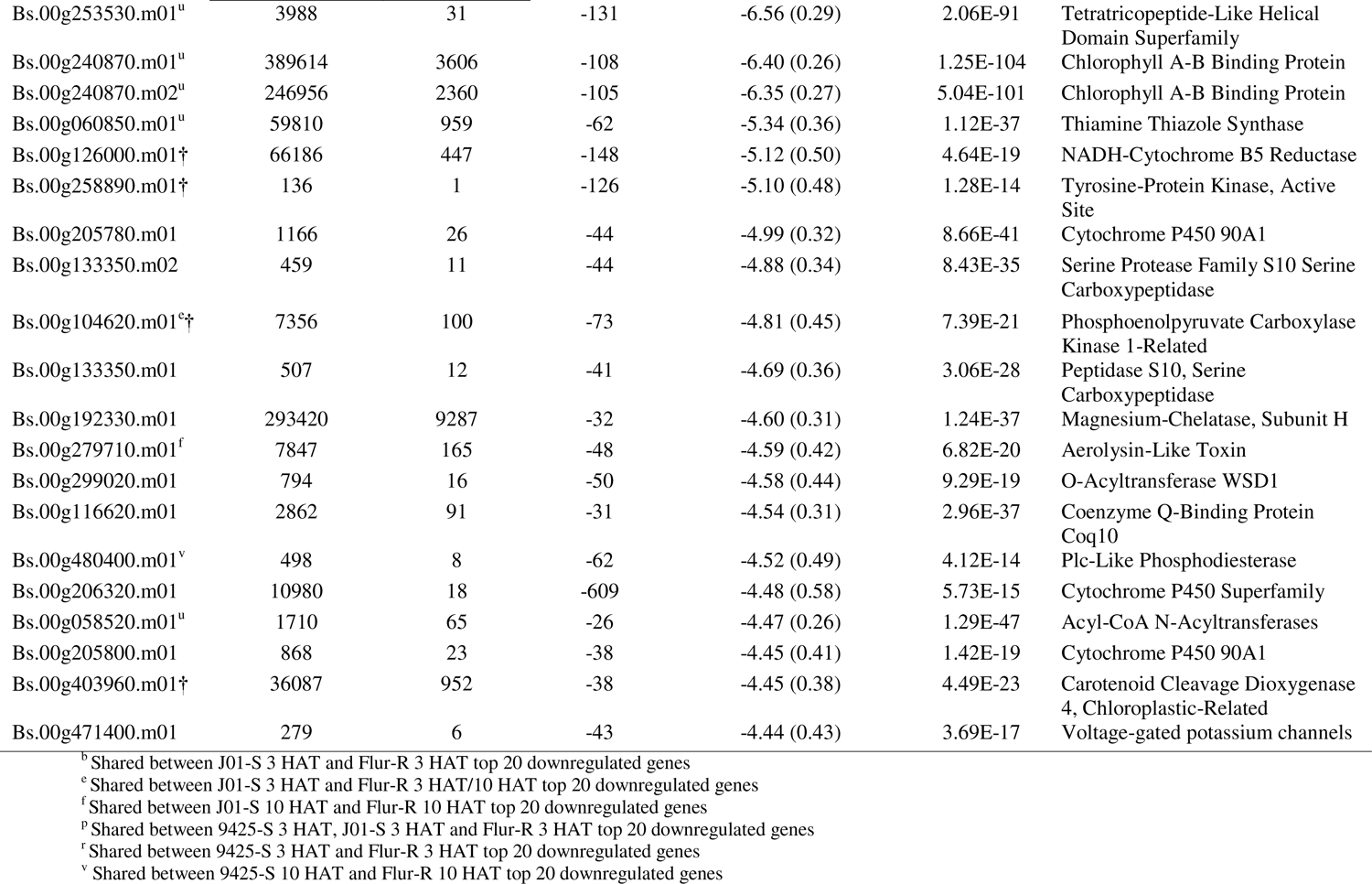
Top 20 downregulated genes in fluroxypyr resistant kochia (*Bassia scoparia*) line Flur-R at 3 h after treatment (HAT) and 10 HAT compared to the untreated timepoint. Fold change was calculated using the mean of normalized counts which was produced using the DESeq2 package in R. Log2 Fold Change was calculated in DESeq2, log2 fold change standard error and adjusted *p*-value were also calculated in DESeq2. The Wald-test obtained *p* values were adjusted using the Benjamini-Hochberg method. The FDR was <0.05.

**Table 7.**
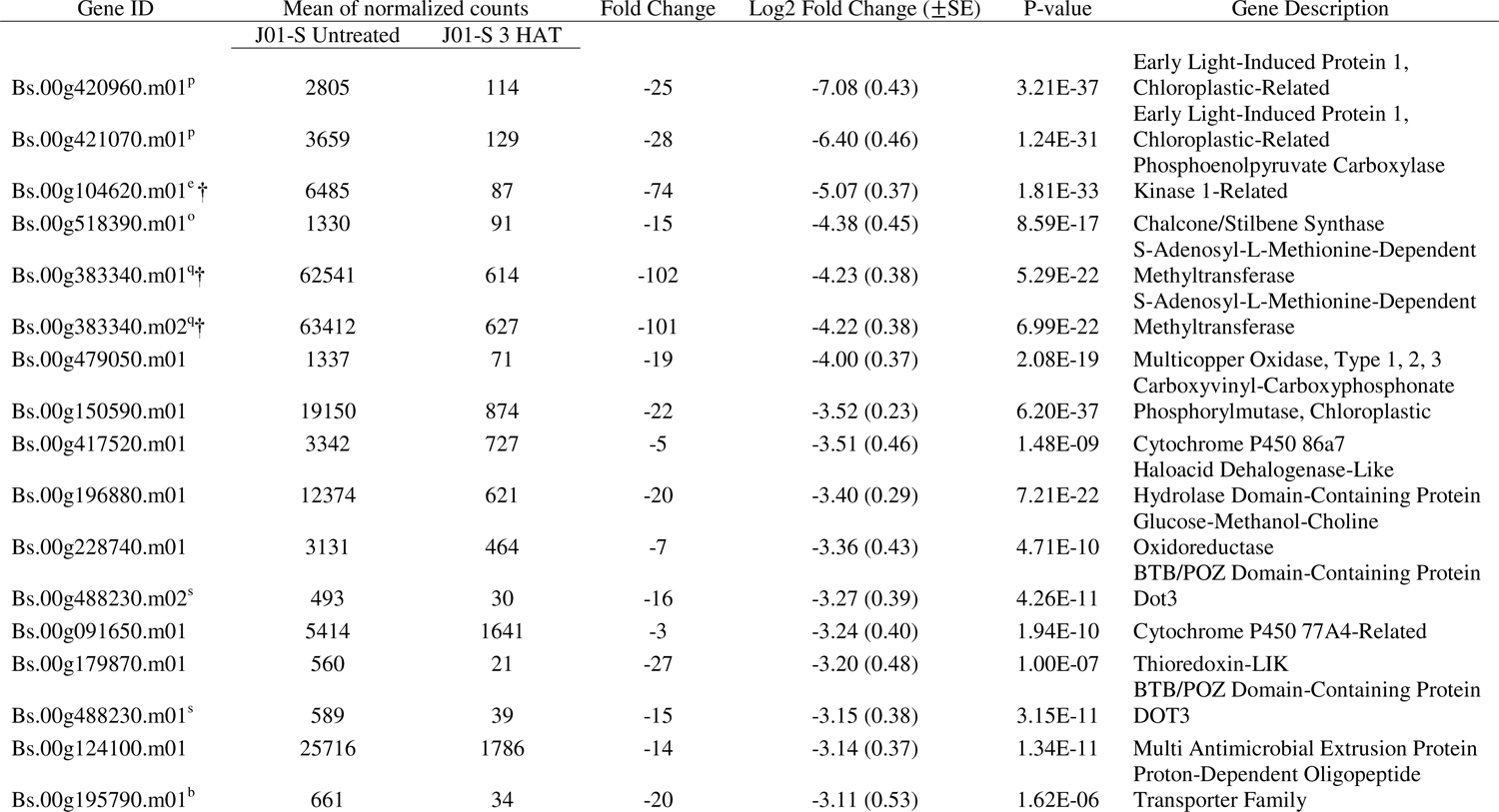
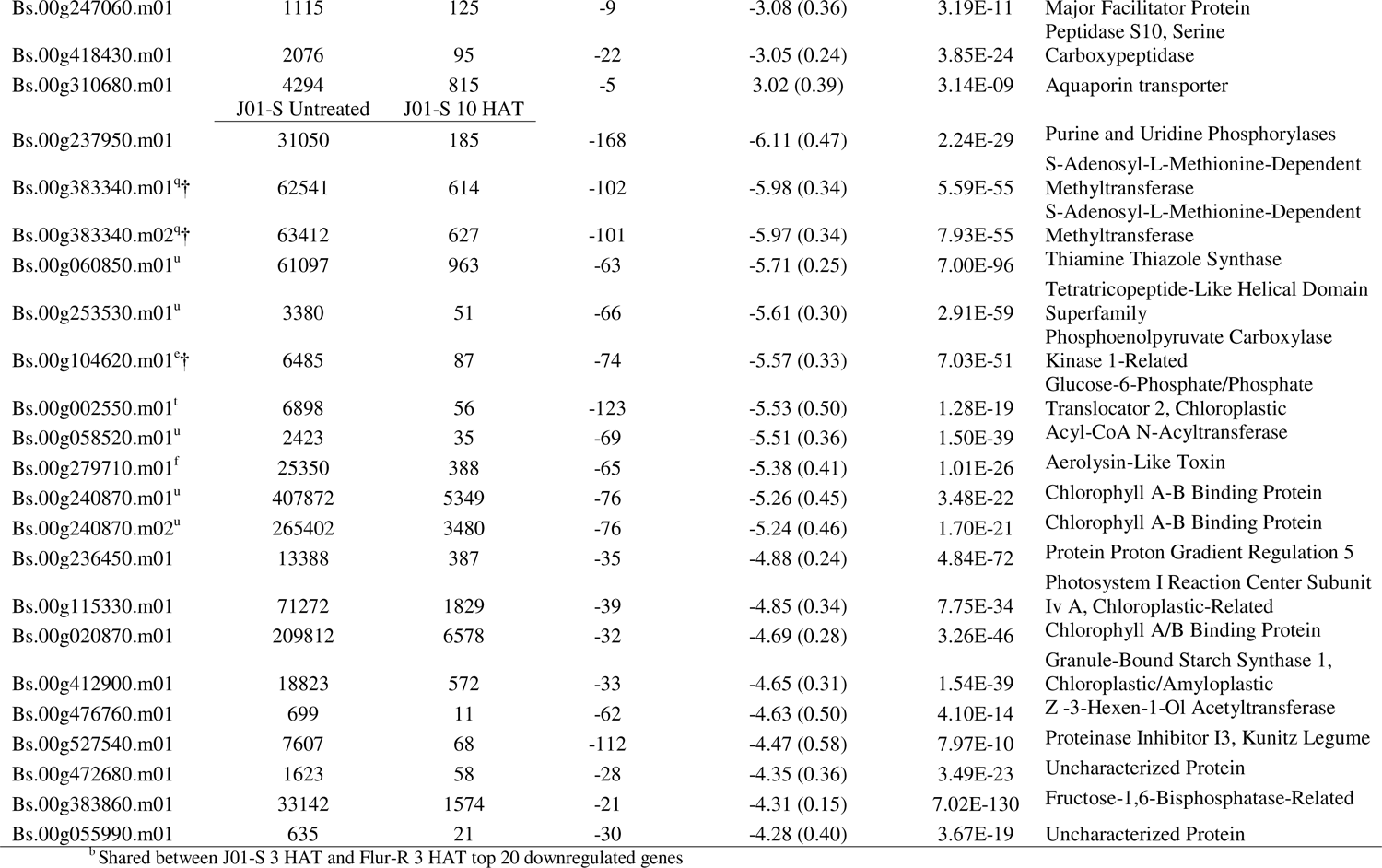
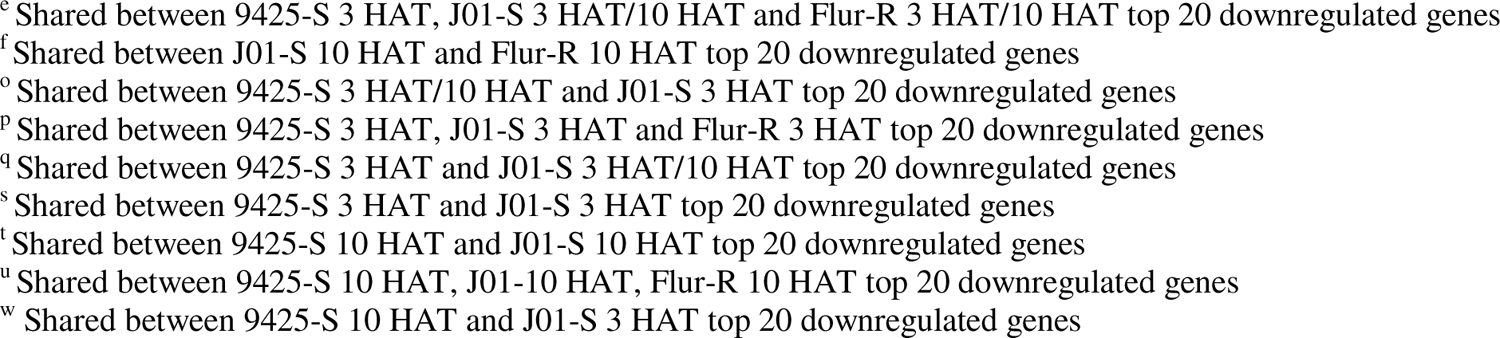
Top 20 downregulated genes in fluroxypyr susceptible kochia (*Bassia scoparia*) line J01-S at 3 h after treatment (HAT) and 10 HAT compared to the untreated timepoint. Fold change was calculated using the mean of normalized counts which was produced using the DESeq2 package in R. Log2 Fold Change was calculated in DESeq2, log2 fold change standard error and adjusted *p*-value were also calculated in DESeq2. The Wald-test obtained *p*-values were adjusted using the Benjamini-Hochberg method. The FDR was <0.05.

**Table 8.**
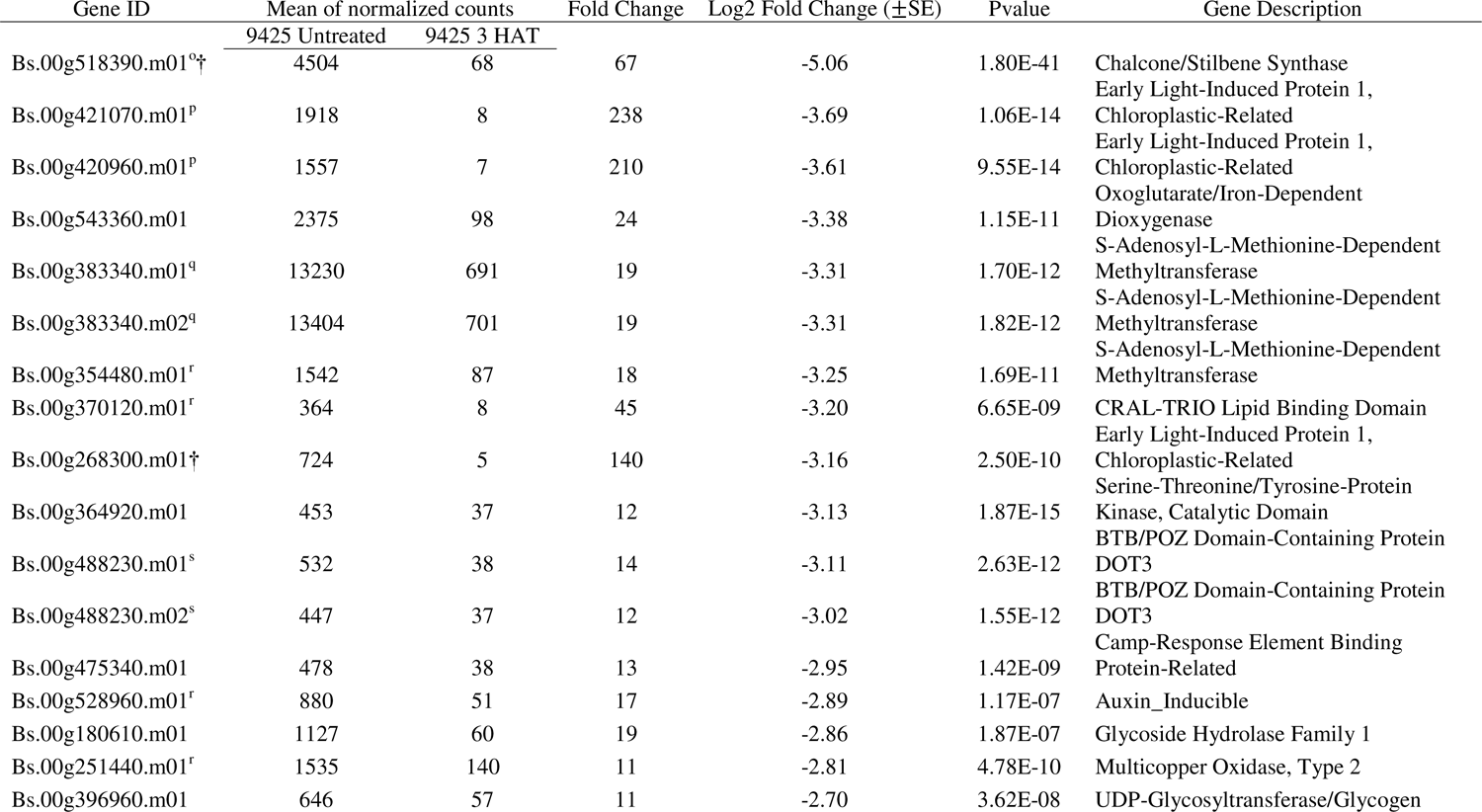
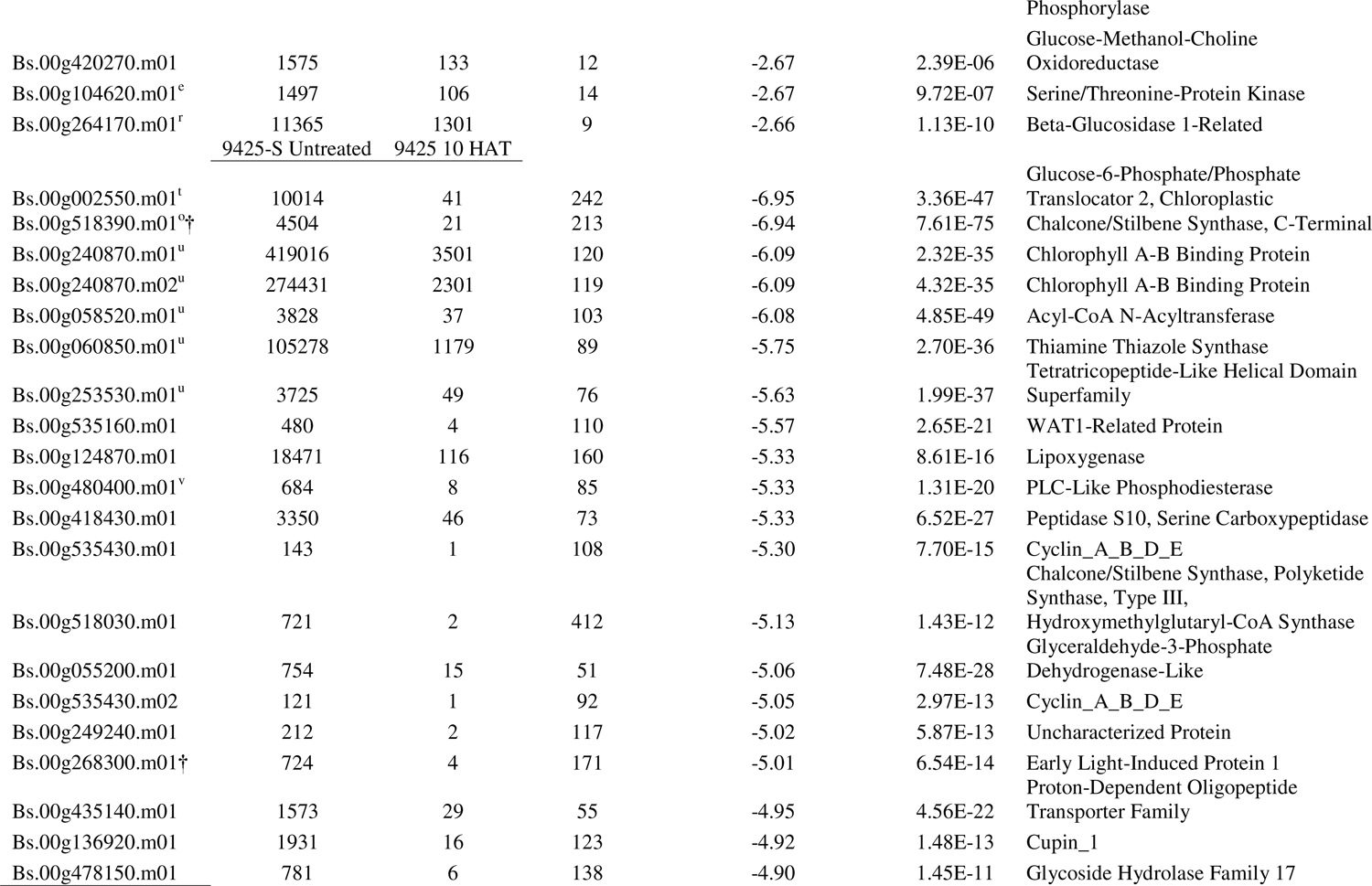
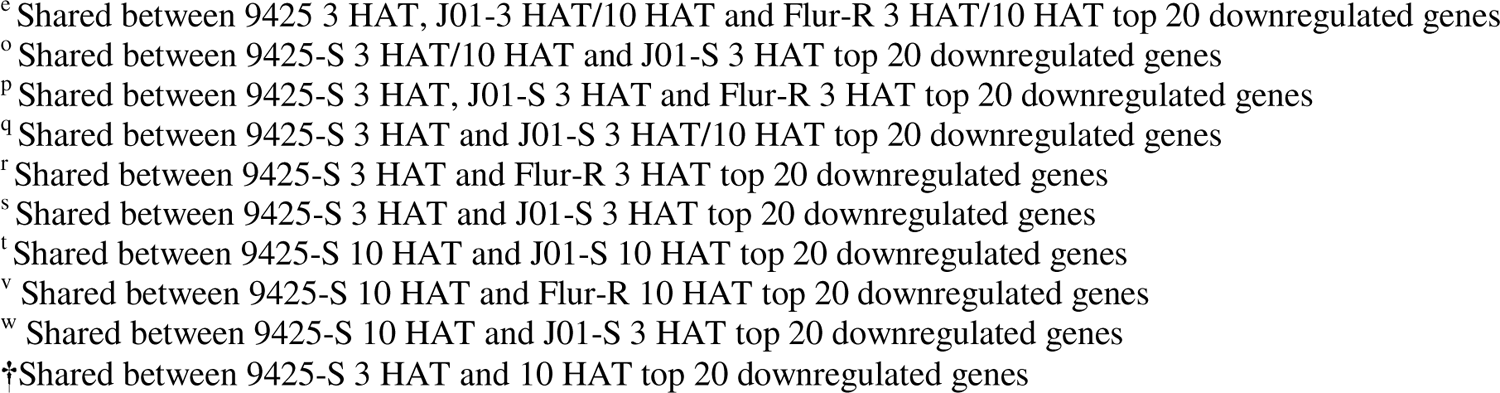
Top 20 downregulated genes in fluroxypyr susceptible kochia (*Bassia scoparia*) line 9425 at 3 hours after treatment (HAT) and 10HAT compared to the untreated timepoint. Fold change was calculated using the mean of normalized counts which was produced using the DESeq2 package in R. Log2 Fold Change was calculated in DESeq2, log2 fold change standard error and adjusted *p*-value were also calculated in DESeq2. The Wald-test obtained *p*-values were adjusted using the Benjamini-Hochberg method. The FDR was <0.05.

### 3.6 Variant Analysis

Our decision criteria to determine resistance-conferring sequence variant candidates were that all four Flur-R individuals from the RNA-seq experiment must have the variant. The candidate variant also must be absent in the two S lines (9425 and J01-S). Of the 147 genes annotated as CYP450s in the kochia genome, there were no unique variants in the Flur-R line. There were 37 genes annotated as having an Aux/IAA domain or function, and 21 genes with an ARF domain or function. Of these genes, three genes contained a nonsynonymous mutation or a deletion. ARF19/7 (*Bs.00g057730.m01*), one of five transcriptional activators in the ARF family, had two nonsynonymous mutations (Gly446Ser; Leu486Ile) and two single codon deletions (SI Figure 3A). A protein annotated as ARF3 (*Bs.00g076170.m01*) also known as ETTIN (ETT) showed one nonsynonymous homozygous variant (Leu293Ser), where three J01-S individuals were heterozygous for the variant found in Flur-R, one was homozygous, and the remaining four 9425 were wildtype. *Aux/IAA4-like* (*Bs.00g107340.m01*) displayed one nonsynonymous variant (Glu52Arg) in a non-conserved region 6-10 bases N terminal of the Aux/IAA Domain II described by Ramos et al. (2001) (SI Figure 3B). We also determined there were no variants in any proteins annotated as AFB or TIR1 proteins that were unique to Flur-R and met our specified criteria, and there were no mutations in the 18 LRR rich C terminus where Aux/IAA and auxin are reported to bind (Villalobos et al. 2012).

## 4. Discussion and Conclusion

Both fluroxypyr resistant line Flur-R and two susceptible lines 9425 and J01-S had up-regulation of auxin regulated genes in response to fluroxypyr that was similar to the auxin mimic herbicide gene expression response in Arabidopsis (Gleason et al. 2011; Goda et al. 2004). The increased expression of auxin responsive genes following fluroxypyr treatment suggests that fluroxypyr is being perceived similarly by all three lines and supports our findings that target-site variants found in *Aux/IAA4* and *ARF19/7* are likely not the cause of the fluroxypyr resistance response in Flur-R. Specifically in *ARF19/7*, the identified variants are predicted to have no significant effect on fluroxypyr binding due to their position in the variable middle region described by Ulmasov et al. (1999) (SI Figure 3A). Although we did find a Flur-R homozygous variant in *ARF3*, the region boundaries of *ARF3* are unlike most other ARFs in that it does not contain Domain III/IV, two key domains for interaction with Aux/IAA proteins relating to auxin gene expression. ARF3 does function in some auxin related pathways (reviewed by Liu et al. (2014)) but the protein has been described to function as a repressor of several proteins causing inhibition of cytokinin activity, a plant hormone that often partners with IAA (Zhang et al. 2018). While we cannot be certain that the variant in *ARF3* does not contribute indirectly to fluroxypyr resistance or affect cytokinin levels in the plant, due to the ARF3 described function, there is stronger support for metabolism being the underlying cause of resistance. Additionally, if variants were found that affected auxin-mimic perception or binding, such as the *IAA16* Gly127Asn mutation described by LeClere et al. (2018), the expected auxin-response gene expression would likely not be induced as reported by Pettinga et al. (2018) in the 9425 line when tested with the auxin-mimic herbicide dicamba.

The translocation data suggest that fluroxypyr, being primarily in its acid form based on the 6 h metabolism results, is moving symplastically throughout the plant as a phloem mobile herbicide (Schober et al. 1986). Transcripts for two IAA transporter (PIN) isoforms were upregulated in the susceptible lines 9425 and J01-S when treated with fluroxypyr, suggesting that PINs can transport fluroxypyr in a similar manner to the transport of IAA. Based on the lack of differences in translocation between Flur-R and J01-S, these two identified PIN transporters are not moving phytotoxic fluroxypyr acid throughout the resistant or susceptible plants at a different rate. Other transporters such as ATP binding cassettes (ABCs) in class B can move multiple substrates including xenobiotics. Some members of this large protein family serve as auxin transporters (Cho and Cho 2013). ABC transporters from both class B and G were upregulated in Flur-R following fluroxypyr treatment, none of which have been individually implicated in herbicide resistance. Several class G transporters are involved in auxin homeostasis and other phytohormone transport, cellular detoxification of heavy metals, and pathogen resistance (Dhara and Raichaudhuri 2021; Gräfe and Schmitt 2021). The functional suite of ABC class G transporters in kochia is yet to be fully described, though cellular export of fluroxypyr conjugates is not outside the scope of known activity for class G transporters.

In Flur-R, abscisic acid (ABA) biosynthesis gene *NCED2* transcript expression decreased over a 10 h time period, contrasting the results from the two susceptible lines in which *NCED6* transcripts had increased expression at 3 h in all three lines (Figure 8). The implications of decreased *NCED2* expression in the resistant line are currently unknown, though some reports show an increased level of response from *NCED* genes following various auxin-mimic applications (Kraft et al. 2007; McCauley et al. 2020; Raghavan et al. 2005). Among these ABA related downregulated genes, seven subunits of Photosystem I and four subunits of Photosystem II are downregulated in all three lines following fluroxypyr application suggesting that fluroxypyr may affect light energy harvesting as part of its mechanism of action. These findings are consistent with the findings of McCauley et al. (2020).

Of the five CYP450s constitutively expressed in Flur-R compared to either 9425 or J01-S, *CYP71D10/11* has been implicated in metabolic herbicide resistance to fenoxaprop-p-ethyl (Bai et al. 2020). Other CYP450s in the CYP71 family have been described as shikimate and shikimate intermediate modifiers (Jun et al. 2015), including Ent-kaurene oxidase (CYP701 subfamily) which functions in gibberellin biosynthesis; its overexpression causes partial resistance to plant growth retardant uniconazole-P (Miyazaki et al. 2011). *CYP81B2* (*Bs.00g431990.m01*) in transgenic tobacco metabolized the phenylurea herbicide chlortoluron after the application of auxin-mimic 2,4-D. The same study also identified its involvement in secondary metabolite biosynthesis (Ohkawa et al. 1999). The other two treatment induced CYP450s in Flur-R, *CYP82D47* and *CYP71A9-like*, have no described role in herbicide resistance, however, there are a significant number of CYP450s involved in herbicide metabolism in the CYP71 family, to which they both belong (Gion et al. 2014; Siminszky et al. 1999; Xiang et al. 2006).

The final constitutively expressed CYP450 in the Flur-R line, *CYP90C1/D1*, belongs to the CYP85 family which is implicated in modification of cyclic terpenes and sterols in brassinosteroid, abscisic acid and gibberellin biosynthesis (Jun et al. 2015; Ohnishi et al. 2006; Ohnishi et al. 2012). It is not unusual for CYP450s to be multifunctional (Bernhardt 2006), and their function can often be attributed to the selectivity of some herbicides, extensively reviewed by Dimaano and Iwakami (2021).

We investigated fluroxypyr resistance using herbicide physiology experiments as well as RNA-sequencing and identified metabolic detoxification as a plausible explanation of fluroxypyr resistance in kochia line Flur-R. Two of the four metabolites are accounted for, having been reported by the Environmental Protection Agency (EPA 2010). The action of conjugation by GSTs or GTs may explain one of the two remaining undescribed metabolites presented, which were both rapidly converted from fluroxypyr acid throughout the time course in Flur-R. Given the high expression of five GSTs and eight GTs in both untreated and treatment induced conditions, formation of secondary metabolic structures is possible. Following CYP450 activity via *O*-glucosylation, fluroxypyr-tripeptide GST or -sugar conjugates can be catalyzed by GST or UDP-glucosyl transferase (Ludwig-Müller 2011). GSTs and GTs can glycosylate plant hormones and xenobiotics to influence bioactivity, transport, solubility and can be pumped out of the cell via ABC transporters (Li et al. 2001; Moons 2005). Subsequent sequestration of the non-phytotoxic herbicide via ABC transporter may also play a role in the resistance response in Flur-R, though more work is needed to fully understand the metabolic response following fluroxypyr application in Flur-R.

## 5. Future Work

Future work elucidating the fluroxypyr resistance mechanism involves *in vitro* and *in vivo* testing of the five candidate GSTs, eight GTs, and eight CYP450s. Metabolite identification via LCMS/MS is crucial next step to determine the metabolic path of the fluroxypyr molecule. Other future studies include genetic mapping of fluroxypyr resistance via test crosses and either Quantitative Trail Loci (QTL) or bulk-segregant analysis with resistant Flur-R and susceptible J01-S, which will provide chromosomal location of resistance gene(s) (Montgomery et al. 2023). Biochemical studies using P450 and GST inhibitors will indicate whether the enhanced fluroxypyr metabolism can be reversed. Metabolic information paired with mapping and ongoing inheritance studies will be a strong contribution to the understanding of auxin-mimic resistance and characterization of fluroxypyr resistance in this population of kochia. Identifying causal resistance genes in auxin-mimic resistant kochia populations will allow us to document the evolution of new resistance genes and predict patterns of gene flow, following the model set by Ravet et al. (2021) for gene flow of glyphosate resistance in kochia.

## Data Availability

The data underlying this article are available in the Gene Expression Omnibus at https://www.ncbi.nlm.nih.gov/geo/query/acc.cgi?acc=GSE179578, and can be accessed with GEO Accession GSE179578.

## Author Contributions Using CRediT Author Statements

**OT:** Writing, visualization, data curation, investigation, formal analysis, validation, methodology **EP:** Resources, methodology **EW:** Resources **AA:** Investigation **WK:** Investigation **FD:** Methodology, supervision, resources **SN:** Methodology, resources **TG:** Writing, supervision, conceptualization.

## Acknowledgements

This research was supported in part by the Colorado Wheat Administrative Committee, by Corteva Agrisciences, and by the USDA National Institute of Food and Agriculture, Hatch project COL00783 to the Colorado State University Agricultural Experiment Station.

