## Supplementary material for "Enhanced metabolic detoxification is associated with fluroxypyr resistance in *Bassia scoparia*": SI Table 1

SI Table 1. Summary of RNA-Seq reads and mapping results. Total read counts, total unmapped reads, uniquely mapped reads and multimapped reads following BGI-seq on fluroxypyr resistant kochia population Flur-R and two fluroxypyr susceptible populations, J01-S and 9425. Three treatments in the RNA-seq study included untreated, “UNT”; 3 hours after treatment with fluroxypyr, “3HAT”; and 10 hours after treatment with fluroxypyr, “10HAT”. Results are taken from the summary text resulting from an alignment of reads to the coding sequence by HISAT2 and feature assignment using featureCounts. Total reads are calculated from the number of pairs. Uniquely mapped reads are calculated from the number of neither concordant nor discordant aligning pairs. Multimapped reads are calculated from the number of concordant pairs that aligned more than once to a given area.

| Sample ID | Total Reads | Unmapped Reads | % Unmapped Reads | Uniquely Mapped Reads | % Uniquely Mapped Reads | Multimapped Reads | % Multimapped Reads | % Alignment |
| --- | --- | --- | --- | --- | --- | --- | --- | --- |
| Flur-R-UNT-1 | 97689782 | 45858498 | 46.94 | 46882608 | 47.99 | 4481566 | 4.59 | 63.75 |
| Flur-R-UNT-2 | 95695742 | 45861354 | 47.92 | 45246428 | 47.28 | 4144868 | 4.33 | 63.23 |
| Flur-R-UNT-3 | 97519984 | 45970566 | 47.14 | 46406360 | 47.59 | 4633858 | 4.75 | 63.61 |
| Flur-R-3HAT-1 | 95413504 | 45414436 | 47.60 | 46002926 | 48.21 | 3546352 | 3.72 | 62.77 |
| Flur-R-3HAT-2 | 91184906 | 43299420 | 47.49 | 43985586 | 48.24 | 3486156 | 3.82 | 63.09 |
| Flur-R-3HAT-3 | 95425988 | 46448452 | 48.67 | 44617614 | 46.76 | 3940164 | 4.13 | 61.85 |
| Flur-R-10HAT-1 | 93693734 | 45778908 | 48.86 | 43823698 | 46.77 | 3675860 | 3.92 | 60.84 |
| Flur-R-10HAT-2 | 93129972 | 46437846 | 49.86 | 42670796 | 45.82 | 3554876 | 3.82 | 60.04 |
| Flur-R-10HAT-3 | 95618080 | 47186278 | 49.35 | 44128110 | 46.15 | 3826944 | 4.00 | 60.99 |
| Flur-R-10HAT-4 | 93585040 | 46051500 | 49.21 | 43362892 | 46.34 | 3764776 | 4.02 | 60.93 |
| 9425-UNT-1 | 92544840 | 47348388 | 51.16 | 41298246 | 44.63 | 3534366 | 3.82 | 60.09 |
| 9425-UNT-2 | 94427486 | 44399484 | 47.02 | 45254046 | 47.92 | 4286030 | 4.54 | 63.2 |
| 9425-UNT-3 | 93239008 | 44049906 | 47.24 | 44570942 | 47.80 | 4187108 | 4.49 | 63.46 |
| 9425-3HAT-1 | 94771882 | 48492414 | 51.17 | 42537404 | 44.88 | 3331316 | 3.52 | 59.72 |
| 9425-3HAT-2 | 94873922 | 49298324 | 51.96 | 41903046 | 44.17 | 3278096 | 3.46 | 58.78 |
| 9425-3HAT-3 | 95517488 | 46581770 | 48.77 | 44650670 | 46.75 | 3815516 | 3.99 | 61.89 |
| 9425-10HAT-1 | 94999072 | 47433958 | 49.93 | 43022690 | 45.29 | 4108722 | 4.33 | 60.68 |
| 9425-10HAT-2 | 94852474 | 48913844 | 51.57 | 41679738 | 43.94 | 3791352 | 4.00 | 59.01 |
| 9425-10HAT-3 | 94430284 | 47108688 | 49.89 | 42689258 | 45.21 | 4167778 | 4.41 | 60.12 |
| 9425-10HAT-4 | 93345484 | 46996468 | 50.35 | 41932346 | 44.92 | 3991146 | 4.28 | 59.94 |
| J01-UNT-1 | 94110422 | 45996468 | 48.87 | 44077502 | 46.84 | 4036452 | 4.29 | 62.26 |
| J01-UNT-2 | 94684556 | 43865012 | 46.33 | 46031172 | 48.62 | 4303572 | 4.55 | 64.28 |
| J01-3HAT-1 | 93575648 | 48396434 | 51.72 | 41217110 | 44.05 | 3496888 | 3.74 | 59.14 |
| J01-3HAT-2 | 94616826 | 46806866 | 49.47 | 43718792 | 46.21 | 3618904 | 3.82 | 61.18 |
| J01-10HAT-1 | 94437294 | 45321298 | 47.99 | 44828044 | 47.47 | 3852384 | 4.08 | 61.64 |
| J01-10HAT-2 | 94714112 | 47245290 | 49.88 | 42739518 | 45.12 | 4292828 | 4.53 | 60.34 |
| J01-10HAT-3 | 94407492 | 46128594 | 48.86 | 43538298 | 46.12 | 4283476 | 4.54 | 61.23 |
