## Supplementary material for "Enhanced metabolic detoxification is associated with fluroxypyr resistance in *Bassia scoparia*": SI Figure 1

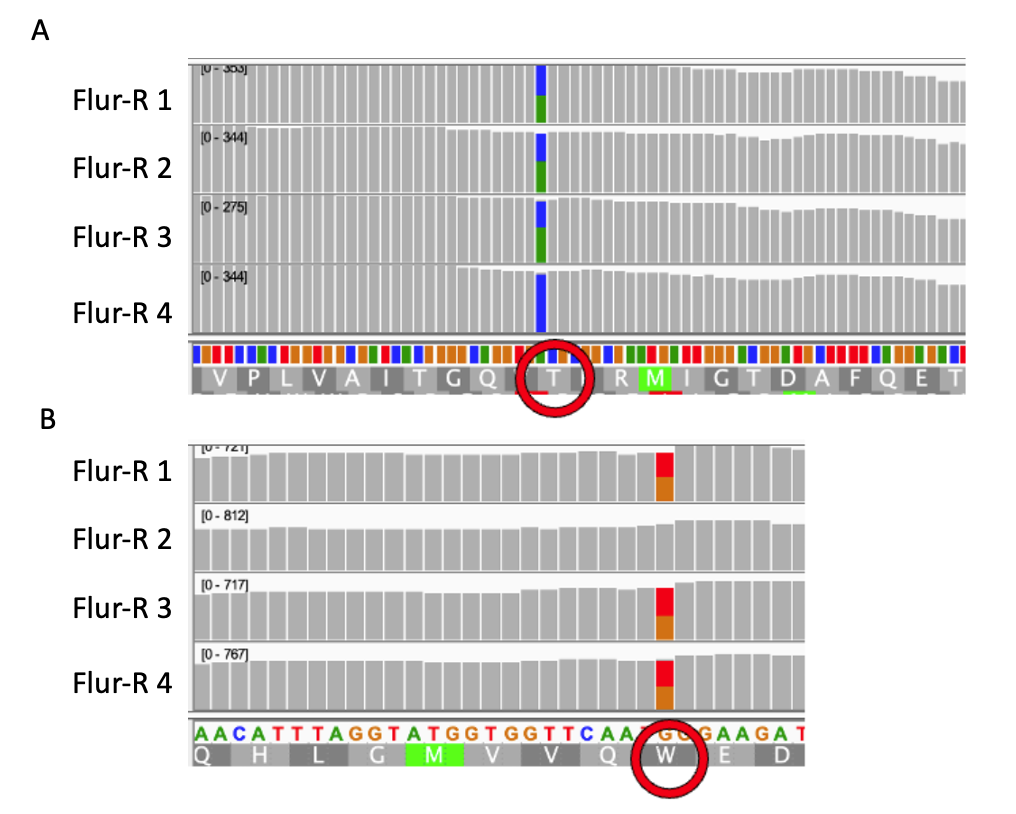

SI Figure 1. Sequence analysis of four Flur-R individuals from the RNA-sequencing data show that there is a Proline 197 to Threonine (P197T) mutation and a Tryptophan 574 to Leucine (W574L) mutation in the acetolactate synthase gene. Resistance to ALS herbicides due to these mutations is a dominant or semi-dominant trait, in which heterozygosity is sufficient to confer resistance to ALS inhibiting herbicides.
