## Supplementary material for "Enhanced metabolic detoxification is associated with fluroxypyr resistance in *Bassia scoparia*": SI Figure 2

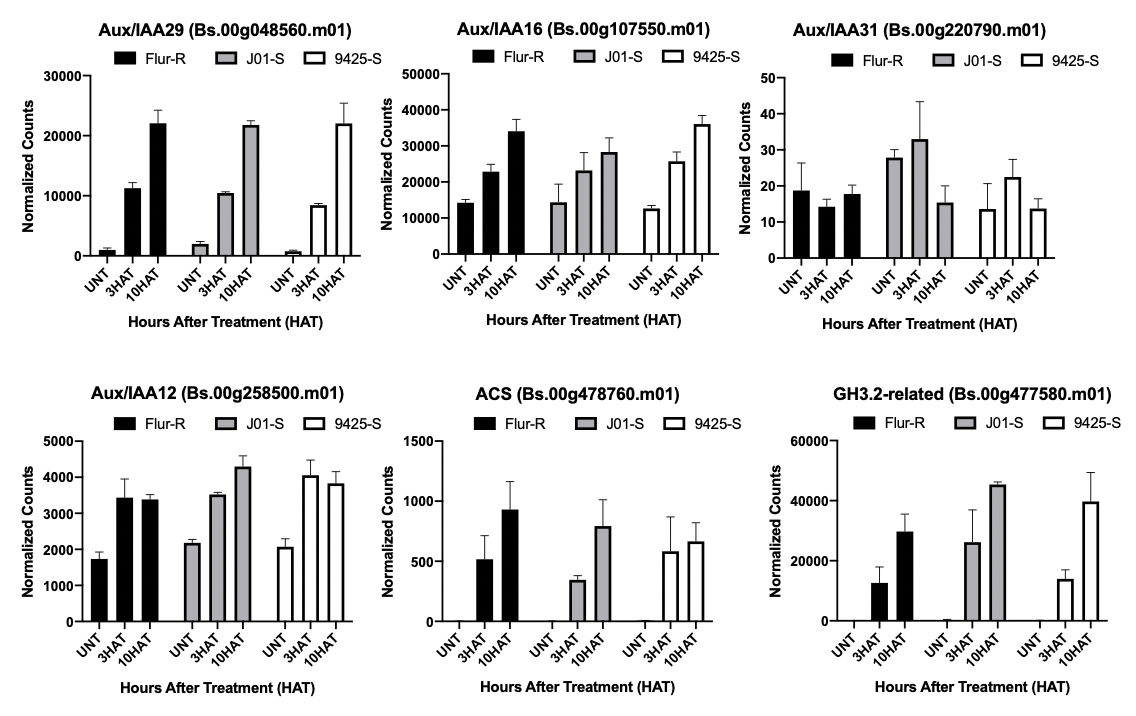
SI Figure 2. Expression profiles for auxin induced genes GH3.2, ACS, and various high annotation confidence Aux/IAAs in fluroxypyr resistant kochia (*Bassia scoparia*) Flur-R, susceptible J01-S, and susceptible 9425-S following differential expression analysis of RNA-Seq data. X-axis shows treatments: untreated, 3 h after treatment (HAT), and 10 HAT grouped by kochia line. Normalized counts on the y-axis are a result of the DESeq2 function and model fitting in R package “DESeq2”.
