## Supplementary material for "Enhanced metabolic detoxification is associated with fluroxypyr resistance in *Bassia scoparia*": SI Figure 3

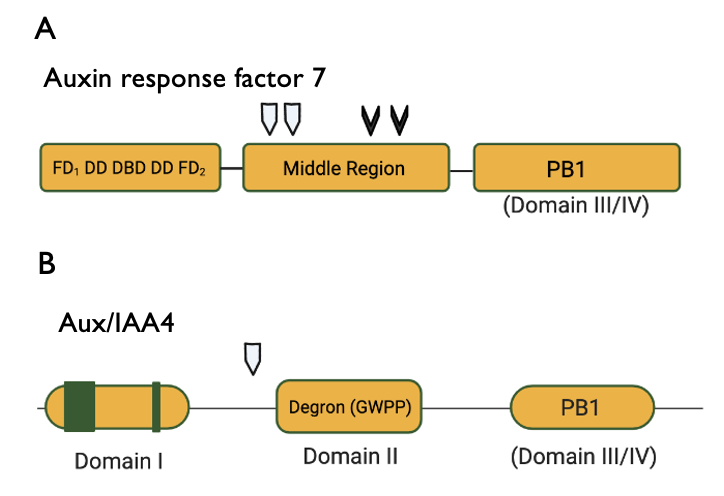

SI Figure 3. A. Variants in the gene ARF 19/7 (Bs.00g057730.m01). Two nonsynonymous mutations (Gly446Ser; Leu486Ile) are represented by white markers, and two single codon deletions are represented by black markers. B. Aux/IAA 4 (Bs.00g107340.m01) nonsynonymous mutation (Glu52Arg) in the N terminal region of Domain II present in the Flur-R line.
